# Cell type composition and circuit organization of neocortical radial clones

**DOI:** 10.1101/526681

**Authors:** Cathryn R. Cadwell, Federico Scala, Paul G. Fahey, Dmitry Kobak, Fabian H. Sinz, Per Johnsson, Shuang Li, R. James Cotton, Rickard Sandberg, Philipp Berens, Xiaolong Jiang, Andreas S. Tolias

## Abstract

**Summary:** Excitatory neurons arising from a common progenitor establish radially-oriented clonal units in the neocortex which have been proposed to serve as elementary information processing modules. To characterize the cell types and circuit diagram within these clonal units, we performed single-cell RNA-sequencing and multi-cell patch clamp recordings of neurons derived from *Nestin*-positive progenitors. We found that radial clones do not appear to be fate-restricted, but instead individual clones are composed of a random sampling of the transcriptomic cell types present in a particular cortical area. The effect of lineage on synaptic connectivity depends on the type of connection tested: pairs of clonally related neurons were more likely to be connected vertically, across cortical layers, but not laterally within the same layer, compared to unrelated pairs. We propose that integration of vertical input from related neurons with lateral input from unrelated neurons may represent a developmentally programmed motif for assembling neocortical circuits.

## Introduction

The mammalian neocortex carries out complex mental processes such as cognition and perception through the interaction of billions of neurons connected by trillions of synapses. We are just beginning to understand how networks of neurons become wired together during development to give rise to cortical computations (Polleux et al., 2007). During cortical neurogenesis, which lasts from approximately embryonic day 10.5 (E10.5) through E17.5 in the mouse (Caviness et al., 1995, Takahashi et al., 1996), radial glial cells (RGCs) undergo asymmetric division to generate postmitotic excitatory neurons that migrate radially to populate the cortical plate. Neurogenesis occurs in an inside-out gradient, such that early-born neurons occupy the deep cortical layers and later-born neurons reside in progressively more superficial layers (Angevine and Sidman, 1961, Rakic, 1974, Caviness et al., 1995). An individual RGC gives rise to a radial unit of clonally related excitatory neurons, sometimes referred to as an *ontogenetic column*, spanning cortical layers 2–6 (Torii et al., 2009, Kriegstein and Noctor, 2004, Noctor et al., 2001, Noctor et al., 2007). However, these radial units of clonally related neurons are only loosely clustered and are intermixed with numerous nearby unrelated neurons (Walsh and Cepko, 1988; Tan et al., 1995). In contrast to excitatory neurons, inhibitory interneurons are generated in specialized regions of the ventral telencephalon and migrate tangentially to disperse throughout the developing cortical mantle (Letinic et al., 2002, Kriegstein and Noctor, 2004, Tan et al., 1998, Mayer et al., 2015).

Recent advances in single-cell RNA-sequencing technology (Tang et al., 2009, Picelli et al., 2013, Picelli et al., 2014a) have enabled unbiased cell type classification in heterogeneous tissues including the cerebral cortex (Zeisel et al., 2015, Tasic et al., 2016, Tasic et al., 2018). An emerging principle is that, in contrast to inhibitory interneurons, excitatory neurons in the adult mouse (Tasic et al., 2018) and developing human (Nowakowski et al., 2017) cortex are largely region-specific at the level of transcriptomic cell types, with several dozens of excitatory cell types per area (Tasic et al., 2018, Hodge et al., 2018). While it is well-established that the vast majority of cells within radial clones are excitatory neurons (Tan et al., 1998), it remains controversial whether individual RGCs can give rise to the full diversity of excitatory neuron cell types within a given cortical area, or whether individual progenitors give rise to a restricted subset of transcriptomic cell types (Eckler et al., 2015, Llorca et al., 2018).

A series of studies used a retroviral lineage tracing method to show that clonally related excitatory neurons are more likely to be synaptically connected to each other (Yu et al., 2009, Yu et al., 2012, He et al., 2015) and also tend to have similar preferred orientations in primary visual cortex (V1) compared to unrelated neurons (Li et al., 2012), consistent with the longstanding hypothesis that radial clones may constitute elementary circuit modules for information processing in the cortex (Mountcastle, 1997, Rakic, 1988, Buxhoeveden and Casanova, 2002). The vertical, across-layer connections between related neurons were described as having a similar directional preference as that found in adult cortex (Yu et al., 2009); however, layer-specific vertical connections were not analyzed independently to directly compare related and unrelated pairs. Moreover, no data were reported for lateral connections between clonally related cells within the same cortical layer. Therefore, it remains unclear whether all local connections are more likely to occur between clonally related neurons, as has become the dogma in the field (Li et al., 2018), or only specific layer-defined connection types. Given the complexity of the local cortical circuit and the different functional roles of layer-defined connections (Lefort et al., 2009, Feldmeyer, 2012, Lubke et al., 2000, Lubke et al., 2003), clarifying the effect of cell lineage on the underlying layer-specific connectivity matrix may have important implications regarding the mechanism and purpose of lineage-driven connectivity. The difficulty of multi-patching experiments combined with the relatively low connectivity rates between excitatory neurons (Jiang et al., 2015, Markram et al., 1997, Barth et al., 2016, Jiang et al., 2016) necessitate testing a very high number of connections and poses an enormous technical challenge to address this question.

Using an enhancer trap Cre-line to label progenitors at an earlier developmental stage, yielding much larger clones (approximately 670-800 neurons per clone compared to 4-6 neurons per clone in Yu et al., 2009), a separate group has reported a much smaller effect of cell lineage on orientation tuning in V1 (Ohtsuki et al., 2012), calling into question the generalizability of cell lineage as an important determinant of large-scale functional circuits (Smith and Fitzpatrick, 2012). While one study has examined lateral connections within layer 4 (L4) of large clones labeled in chimeric mice (Tarusawa et al., 2016), the effect of cell lineage on other layer-defined connection types has not been systematically studied in large clones. This represents a major obstacle in translating the idea of lineage-dependent circuit assembly into a practical model of cortical circuit development.

Here we use a tamoxifen-inducible Cre-lox system to label progenitors precisely at the onset of neurogenesis, resulting in intermediate-sized radial clones (86 neurons on average; Figure 1) spanning cortical layers 2-6 to ask: a) what is the cell type composition of individual clones and b) what is the layer-specific connectivity matrix among clonally related excitatory neurons. We find that radial clones of excitatory neurons are composed of diverse transcriptomic cell types with no evidence of cell type fate restriction compared to nearby unrelated neurons. In addition, vertical connections linking cells across cortical layers, and in particular vertical inputs to cortical layer 5, were selectively increased among clonally related neurons with no change in lateral connections within the same cortical layer. These findings suggest a revision of the current dogma of universally increased connectivity among clonally related excitatory neurons and suggest that integration of vertical input from related neurons within radial units and lateral input from unrelated neurons may represent a developmentally programmed blueprint for the construction of functional neocortical circuits.

**Figure 1.**
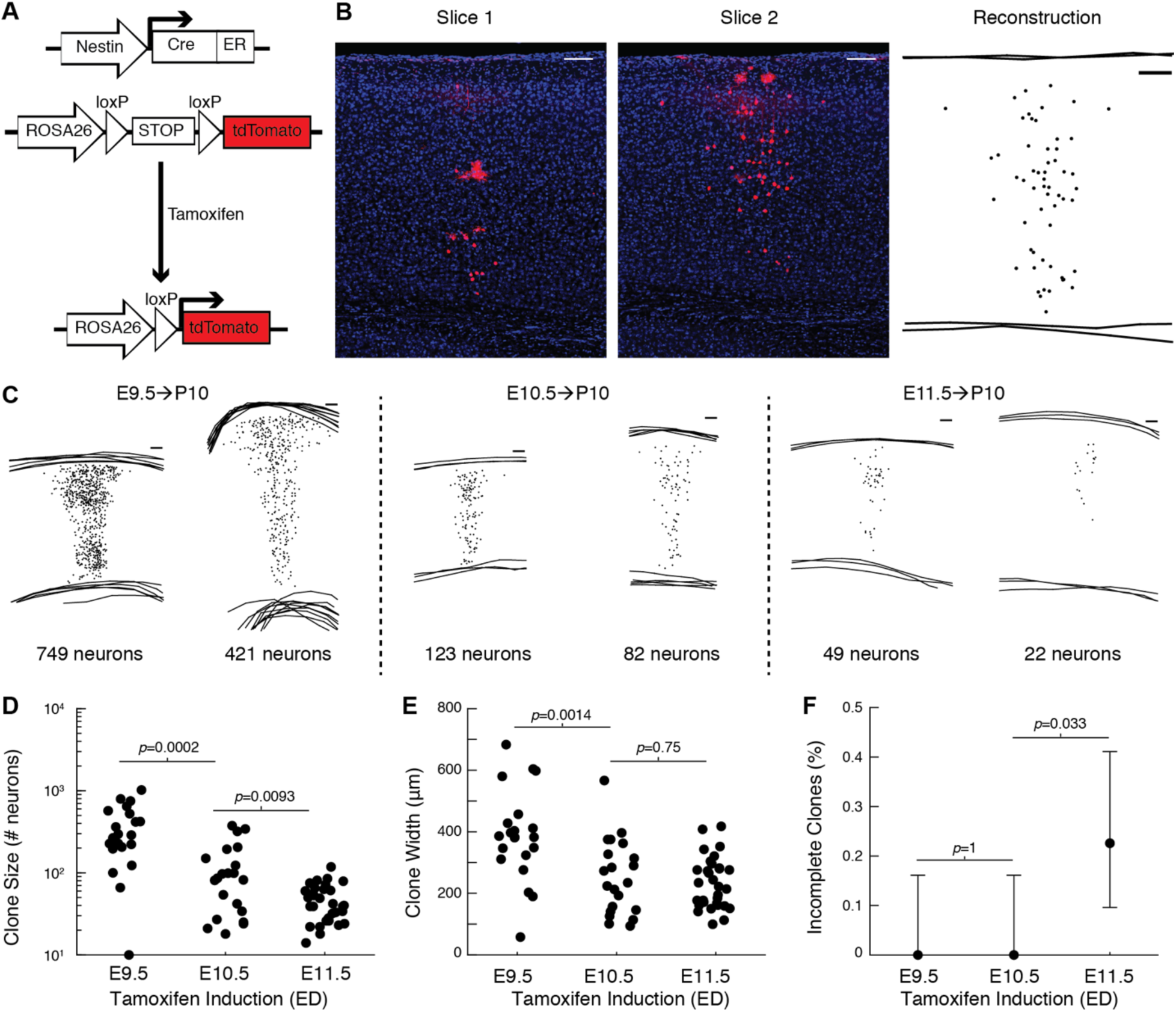
Tamoxifen induction at E10.5 generates radial clones spanning superficial and deep cortical layers. **(A)** Schematic of tamoxifen-inducible Cre-loxP system for lineage tracing. **(B)** Manual reconstruction of clone across multiple slices. In this example, larger red spots are morphologically consistent with glial cells at high magnification. Scale bar: 100 µm. **(C)** Examples of reconstructed clones labeled at E9.5, E10.5 or E11.5. Scale bar: 100 µm. **(D and E)** Number of neurons **(D)** and clone width **(E)** at postnatal day 10 following tamoxifen induction at E9.5, E10.5, or E11.5 (n = 21, 21, and 31 clones; n = 2 mice per condition; *p*-values computed using Wilcoxon rank sum. **(F)** Percent of clones that are incomplete (do not include L5–L6) following tamoxifen induction at E9.5, E10.5, or E11.5 (n=21, 21, and 31 clones; n = 2 mice per condition; *p*-values computed using Fisher’s exact test). Error bars show 95% Clopper-Pearson confidence intervals. See also Figure S1.

## Results

### Tamoxifen induction at E10.5 generates radial clones spanning superficial and deep cortical layers

To label radial clones, we induced sparse recombination in RGCs at approximately the onset of neurogenesis using a tamoxifen-inducible Cre-lox transgenic system driven by the *Nestin* promoter (Figures 1A-1B). In contrast to viral lineage tracing methods, which are routinely performed at E12.5 or later (Yu et al., 2009, Yu et al., 2012, Li et al., 2012), and enhancer trap methods (Ohtsuki et al., 2012), which do not allow precise control of the labeling time (i.e. number of neurons per clone) or density (i.e. numer of clones per brain), our approach enabled us to empirically determine the optimal dosing and time point to sparsely label radial clones (Figure 1C). When tamoxifen was administered shortly before the onset of neurogenesis, at E9.5, the resulting clones at P10 contained 288 neurons (median, interquartile range [IQR] 196–525 neurons) and spanned 383 µm in width (median, IQR 311–428 µm; n=21 clones from two animals; Figures 1D–E). When tamoxifen was administered at approximately the onset of neurogenesis, at E10.5, the resulting clones at P10 contained 86 neurons (median, IQR 27–150) and spanned 235 µm in width (median, IQR 141–327 µm; n=21 clones from two animals; Figures 1D–E). The substantial decrease in clone width and neuron number between clones labeled at E9.5 and those labeled at E10.5 suggests inducing recombination prior to E10.5 leads to the labeling of a substantial number of neuroepithelial stem cells still undergoing symmetric cell division to generate multiple radial glial cells. When tamoxifen was administered shortly after the onset of neurogenesis, at E11.5, the resulting clones at P10 contained 40 neurons (median, IQR 26–61) and spanned 222 µm in width (median, IQR 162–280 µm; n=31 clones from two animals; Figures 1D–E), similar to the width of clones labeled at E10.5. Moreover, a substantial fraction of clones labeled by induction at E11.5 were restricted to the superficial cortical layers (L2–4; 7/31 clones, 23%), which was never seen in clones labeled by induction at E9.5 or E10.5 where they spanned layers 2 to 6 (Figures 1F and S1). This finding is consistent with the model that radial glia contribute to all excitatory cortical layers, but also suggests that at least some radial glia no longer generate deep layer neurons after E11.5, consistent with the inside-out model of excitatory neurogenesis. Given the plateau of clone width when induced at E10.5 or E11.5 and the presence of incomplete clones when induced at E11.5, we reasoned that E10.5 is the optimal time point to induce recombination in order to label radial clones spanning both superficial and deep cortical layers, and we used this induction protocol for all of our subsequent experiments.

### Transcriptomic variability of excitatory neurons is driven primarily by layer position and cortical region

A recent transcriptomic cell atlas of adult mouse neocortex showed that primary visual (V1) and anterior lateral motor (ALM) cortices are composed of distinct transcriptomic excitatory neuron cell types (Tasic et al., 2018). Given that our lineage tracing strategy labels radial clones randomly across many cortical regions, we next asked whether region-specific excitatory neurons are present also in juvenile mice.

To test this, we cut acute parasagittal slices spanning primary visual (V1) and primary somatosensory (S1) cortices from juvenile (P15-P20) mice. Radial clones were identified by their intrinsic fluorescence (Figure 2A–B) and the contents of individual neurons were aspirated through a patch pipette following brief electrophysiological recording using our recently described Patch-seq protocol (Cadwell et al., 2017, Cadwell et al., 2016, Fuzik et al., 2016). We analyzed 206 neurons (after quality control, see Figure S2 and Methods) which had approximately 0.36 million uniquely mapping reads (median; IQR 0.17–0.69 million reads; Figure 2C) and approximately 7000 genes detected (median: 7007; IQR 6152–7920 genes; Figure 2D). For all downstream analyses, we used 12,841 genes that had on average >1 count/cell (see Methods). There were modest differences in the average library size (Figures S2E–F) and number of genes expressed (Figures S2I–J) between different cortical areas and between different layers. However, there were no significant differences between tdTomato-positive (clonally related) and tdTomato-negative (nearby unrelated) neurons in either of these two measures (Figures S2G and S2K). Count data were normalized using a pool-based strategy developed specifically for single-cell RNA sequencing analysis (Lun et al., 2016). Size factors largely correlated with library size (Figure S2H), suggesting that our cell population is relatively homogenous and that systematic differences in gene counts in our dataset are driven primarily by technical factors such as capture efficiency and sequencing depth. The normalized counts (Table S1) were used in all subsequent analyses.

**Figure 2.**
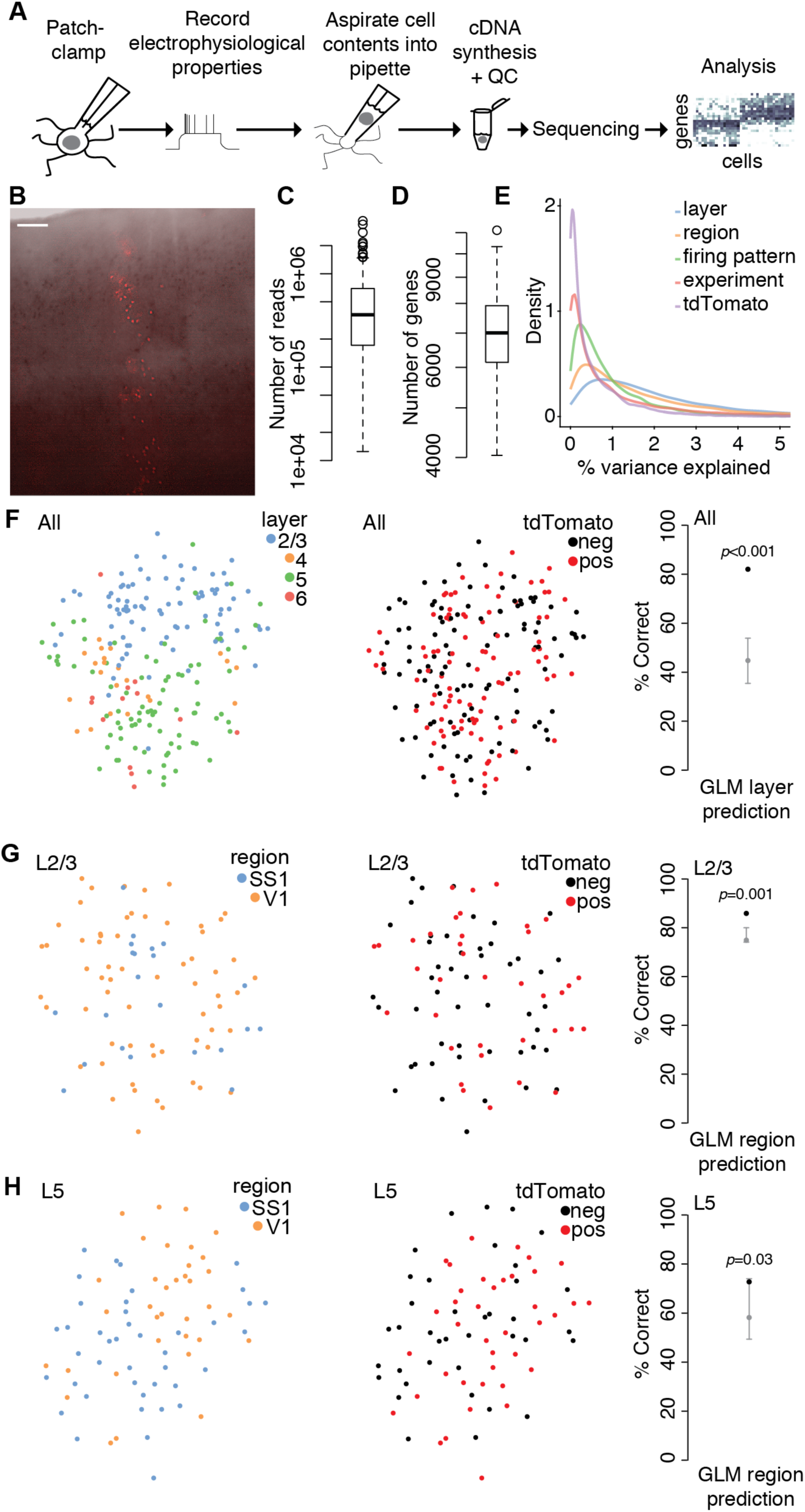
Transcriptomic variability of excitatory neurons is driven primarily by layer position and cortical region. **(A)** Overview of experimental approach using Patch-seq. **(B)** Example tdTomato-positive radial clone spanning superficial and deep cortical layers in an acute cortical slice used for Patch-seq experiments. Overlay of bright field and fluorescence image was performed in Adobe Photoshop. Scale bar: 100 µm. **(C and D)** Box plots showing library size **(C)** and number of genes detected **(D)** for all cells passing quality control criteria (n=206). **(E)** Density plot of the percent of variance in normalized log-expression values explained by different experimental factors across genes (n=12,841 genes). **(F)** T-distributed stochastic neighbor embedding (t-SNE) plots using the top highly variable and correlated genes across all cells (n=91 genes; n=87, 22, 84, and 13 cells in layers 2/3, 4, 5, and 6, respectively), colored by layer position **(left)** or tdTomato expression **(middle)**. **(right)** Performance of a generalized linear model (GLM) trained to predict layer position from gene expression data (n=12,841 genes and 206 cells) with model performance (black dot) compared to the chance-level performance estimated using shuffled data (grey, mean and 95% coverage interval; shuffling layer). one-tailed *p*-value computed from shuffled data. **(G)** t-SNE plots using the top highly variable and correlated genes across L2/3 cells (n=43 genes; n=22 and 63 cells in SS1 and V1, respectively), colored by region **(left)** or tdTomato expression **(middle)**. **(right)** Performance of a GLM trained to predict region from gene expression data (n=12,841 genes and 85 cells) as described in **F** but shuffling region instead of layer. **H)** t-SNE plots using the top highly variable and correlated genes across L5 cells (n=41 genes; n=42 and 35 cells in SS1 and V1, respectively), colored by region **(left)** or tdTomato expression **(middle)**. **(right)** Performance of a GLM trained to predict region from gene expression data (n=12,841 genes and 77 cells) as described in **F** but shuffling region instead of layer. See also Figures S2 and S3 and Table S1.

Consistent with recent studies (Zeisel et al., 2015, Tasic et al., 2016, Tasic et al., 2018, Nowakowski et al., 2017), we found that layer position and cortical region were strong predictors of transcriptomic variability among excitatory neurons in our dataset (Figures 2E and S3). Dimensionality reduction using t-distributed stochastic neighbor embedding (t-SNE) revealed that neurons clustered primarily by layer position, and a cross-validated generalized linear model (GLM) could predict layer position from the gene expression data with approximately 80% accuracy (Figure 2F). Within L2/3 and L5, neurons from V1 and S1 seemed to form overlapping clusters but GLM prediction accuracy was slightly better than chance (Figures 2G and 2H), suggesting that in juvenile mice, L2/3 and L5 excitatory neurons may have already started differentiating into region-specific transcriptomic classes.

### Radial clones are composed of diverse transcriptomic subtypes of excitatory neurons with no evidence of fate restriction

While most evidence supports a deterministic model of excitatory neurogenesis, whereby individual progenitors give rise to many different excitatory neuron cell types through progressive fate restriction (Tan and Breen, 1993, Guo et al., 2013, Gao et al., 2014), other studies suggest that a subset of progenitors may be fate-restricted early on to give rise to layer-restricted excitatory neurons (Franco and Muller, 2013, Franco et al., 2012, Gil-Sanz et al., 2015, Llorca et al., 2018) and the relative contribution of the “common” and “multiple” progenitor models in generating excitatory neuron diversity remains controversial. Moreover, the transcriptomic diversity of clones of excitatory neurons is unknown, and one possibility is that individual radial clones may give rise to only a subset of the various cell types present within a given cortical layer; for example, some have proposed that up to one quarter of radial clones spanning L2-L6 may be composed exclusively of corticocortical projection neurons (Llorca et al., 2018).

To characterize the diversity of cell types within radial clones, we mapped our single-cell transcriptional profiles to a recently published cell type atlas of adult mouse cortex (Tasic et al., 2018). We found that labeled neurons within radial clones mapped to all of the broad excitatory cell classes (Figures 3A and S4A, and Table S2) in proportions similar to the unlabeled control neurons (Figures 3B and S4B), suggesting that the *Nestin*-positive progenitors labeled using our lineage tracing protocol can give rise to the full range of excitatory neuronal cell types in the cortical areas examined. Area S1 was not specifically profiled in the reference cell atlas; we found that cells from both V1 and S1 in our dataset mapped predominantly to V1 excitatory neuron types (92.7% of all cells, n=191/206; 94.0% of V1 cells, n=110/117; 96.2% of S1 cells, n=76/79; Table S2) with only a handful mapping to ALM excitatory neuron types (4.9% of all cells, n=10/206; 6.0% of V1 cells, n=7/117; 3.8% of S1 cells, n=3/79; Table S2). The quality of the mapping was equally good for V1 and S1 cells (mean uncertainty for S1 cells, 5.8±0.9; mean uncertainty for V1 cells, 5.0±0.6; mean±SE in arbitrary units (see Methods); p=0.44, two-sample t-test), suggesting that the adult V1/ALM cell type atlas is an equally reasonable reference for excitatory neurons in juvenile V1 and S1. As a sanity check, we observed that most neurons patched in L2/3 mapped to L2/3 reference types (65/87, 74.7%), and similarly for L4 (15/22, 68.2%), L5 (56/84, 66.7%), and L6 (11/13, 84.6%) (Figure 3A and Tables S1 and S2). The discrepancies were mostly due to some neurons mapping to a transcriptomic type from a neighboring layer: neurons from L2/3 mapping to L4 types (8/87, 9.2%), neurons from L5 mapping to L4 types (5/84, 6.0%), and neurons from L5 mapping to L6 types (19/84, 22.6%).

**Figure 3.**
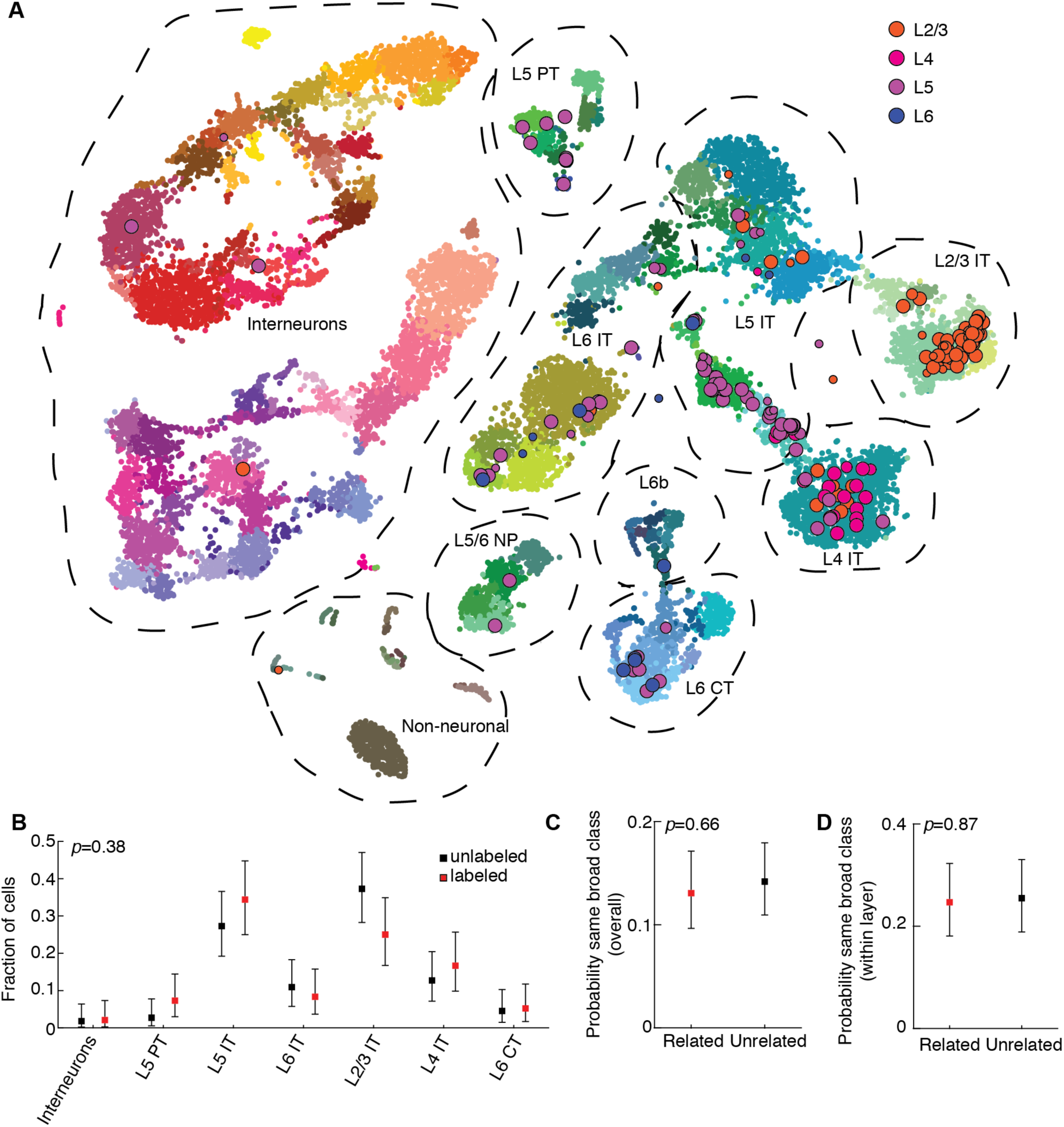
Radial clones are composed of diverse transcriptomic subtypes of excitatory neurons with no evidence of fate restriction. **(A)** T-distributed stochastic neighbor embedding (t-SNE) plot showing alignment of our Patch-seq data (data points with black outline, n=87, 22, 84, and 13 cells in layers 2/3, 4, 5, and 6 respectively) with a recently published mouse cell type atlas (data points with no outline; n=23,822; from (Tasic et al., 2018); colors denote transcriptomic types and are taken from the original publication). The t-SNE of the reference dataset and the positioning of Patch-seq cells were performed as described in (Kobak & Behrens, 2018), see Methods. The size of the Patch-seq data points denotes the precision of the mapping (see Methods): small points indicate high uncertainty. **(B)** Fraction of labeled (n=96) and unlabeled (n=110) cells that mapped to each of the broad cell classes outlined in **(A)** with greater than three Patch-seq cells total. **(C and D)** Probability of related and unrelated cell pairs mapping to the same broad cell class either overall (**C**; n=337 related pairs, n=409 unrelated pairs) or when conditioned on layer position (**D**; n=154 related pairs, n=157 unrelated pairs). For **(B–D)**, error bars are 95% Clopper-Pearson confidence intervals and *p*-values are computed using Chi-squared test. See also Figures S4 and S5 and Table S2.

Within individual radial clones, pairs of related neurons were no more likely to map to the same broad transcriptomic class (Figures 3C 3D and S5), or specific cell type (Figures S4C S4D, and S5) compared to pairs of unrelated neurons. Our results do not support the model that a subset of cortical radial glia are fate-restricted. Instead, our data is consistent with a “single progenitor” model of excitatory neurogenesis in which individual progenitors are capable of generating all of the diverse excitatory neuronal types within a given cortical region.

### Vertical, across-layer connections are selectively increased between excitatory neurons in radial clones

To determine whether clonally related neurons within our radial clones were preferentially connected, we performed multiple simultaneous whole-cell recordings as previously described (Jiang et al., 2015), targeting up to eight neurons simultaneously including both clonally related cells and nearby unlabeled control cells (Figures 4A and 4B). In total, we patched 592 neurons (310 labeled and 282 unlabeled) from 86 clones in 43 mice. The cells were distributed throughout L2/3 (n=275 cells), L4 (n=164 cells) and L5 (n=153 cells). To test connectivity, we injected brief current pulses into each patched neuron to elicit action potentials and monitored the responses of all other simultaneously recorded neurons to identify unitary excitatory postsynaptic currents (uEPSCs, Figure 4C). To confirm that the recorded cells were excitatory neurons, we analyzed the firing pattern of each cell in response to sustained depolarizing current and examined the morphology of each neuron using avidin-biotin-peroxidase staining (see Methods). In addition, we measured the inter-soma distances between all pairs of simultaneously recorded neurons. Cells that did not show definitive electrophysiological and/or morphological features of excitatory neurons (5.9%, 35/592) were excluded from further analysis. In total, we tested 2049 potential excitatory connections and identified 112 synaptic connections. The uEPSCs had a latency of 2.71±1.06 ms (n=112 connections analyzed; mean±SD), an amplitude of 12.83±14.11 pA (n = 112 connections analyzed, mean±SD), and were blocked by bath application of glutamatergic antagonists CNQX (20 µM) and APV (100 µM; uEPSC amplitude = 10.5±5.4 pA and 0.0±0.0 pA before and after the application of antagonists; median±SE; n=15 connections tested, *p*=6×10^−5^, Wilcoxon signed-rank test), further confirming that these were excitatory connections.

**Figure 4.**
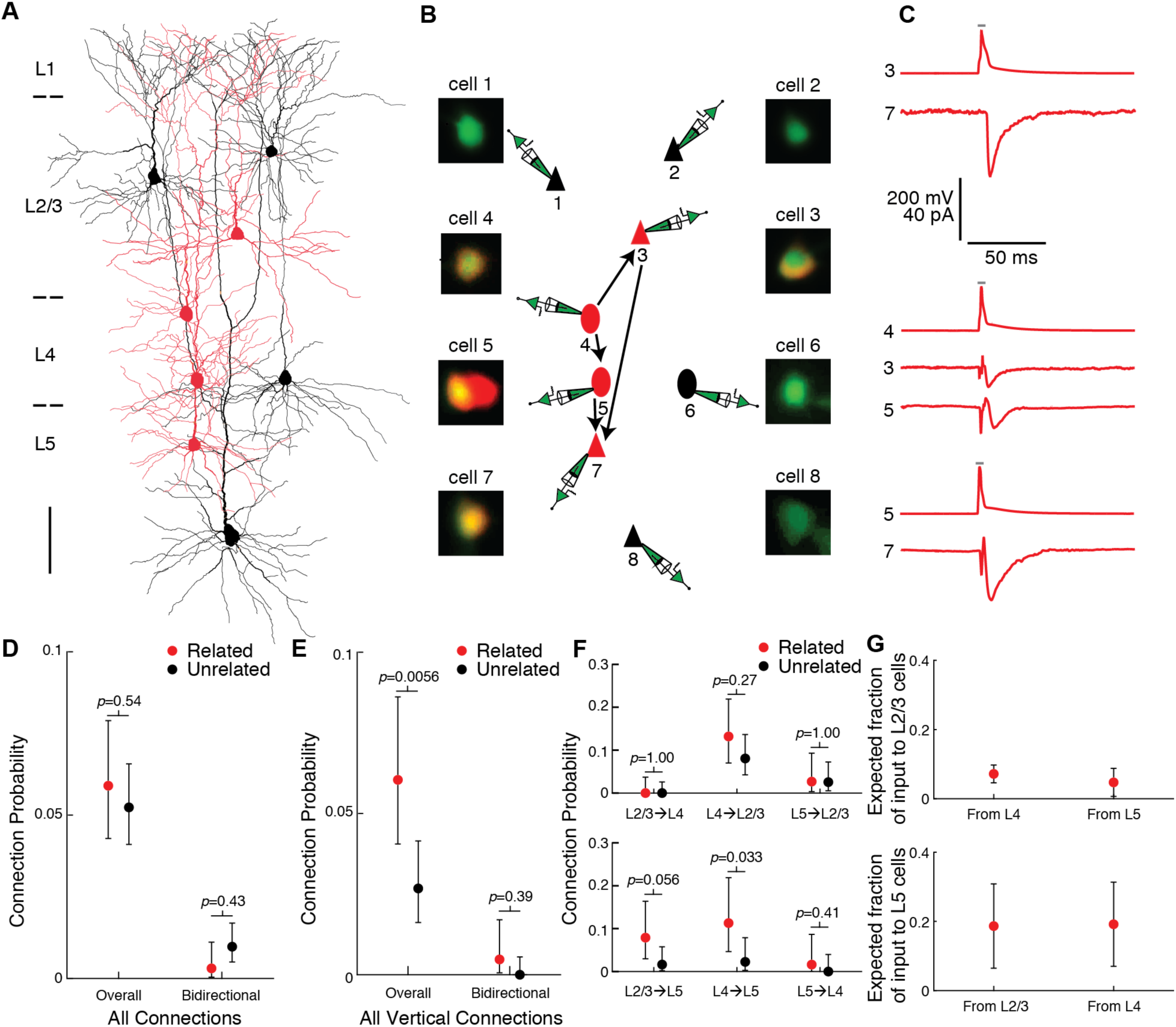
Vertical, across-layer connections are selectively increased between excitatory neurons in radial clones. **(A–C)** Example recording session from four clonally related cells (red) and four nearby, unrelated control cells (black). **(A)** Morphological reconstruction of all eight neurons. Scale bar, 100 µm. **(B)** Schematic of connections identified, as well as fluorescence images of each patched cell confirming the overlap of red (lineage tracer) and green (pipette solution) in related cells and green only in control cells. Triangles, pyramidal neurons; ovals, L4 excitatory neurons. **(C)** Presynaptic action potential (AP) and postsynaptic uEPSC traces for each connection (average of at least 30 trials each). Grey bar indicates period of depolarizing current injection to presynaptic neuron. **(D)** Connection probabilities among related and unrelated neurons, pooling all connections tested (n=712 potential connections and 324 pairs with both directions tested for related neurons; n=1337 potential connections and 612 pairs with both directions tested for unrelated neurons). **(E)** Connection probabilities among related and unrelated neurons, pooling all vertical, across-layer connections tested (n=464 potential connections and 211 pairs with both directions tested for related neurons; n=711 potential connections and 333 pairs with both directions tested for unrelated neurons). **(F)** Connection probabilities among related and unrelated neurons, for each vertical connection type tested (n=98, 91, 75, 76, 62, and 62 potential connections for related neurons and n=141, 149, 118, 123, 89, and 91 potential connections for unrelated neurons from L2/3 to L4, L4 to L2/3, L5 to L2/3, L2/3 to L5, L4 to L5, and L5 to L4, respectively). **(G)** Estimated fraction of vertical, across layer input to L2/3 cells (top panel) and L5 cells (bottom panel) coming from clonally related neurons based on our empirically measured clone sizes and connection probabilities. For **(D–F)**, error bars are 95% Clopper-Pearson confidence intervals and *p*-values are computed using Fisher’s exact test. For **(G)**, error bars are propagated standard error of the estimates (see Methods). See also Figures S7 and S8.

To determine the effect of cell lineage on connection probability (P), we compared pairs consisting of two labeled cells within the same clone (i.e. “related” pairs) to pairs consisting of one labeled and one unlabeled cell (i.e. “unrelated” pairs). Pairs consisting of two unlabeled cells were not included as controls, since we could not be certain that those pairs are unrelated (they could be related, but their progenitor was not labeled). Overall, there was no evidence for a difference in connectivity between related and unrelated pairs (P=5.9% [42 out of 712 potential connections] and P=5.2% [70 out of 1337] for related and unrelated pairs, respectively; *p*=0.54, Fisher’s exact test; Figure 4D). A single bidirectional connection was identified between related neurons (0.0031%, 1 out of 324 pairs in which both directions of connectivity were tested), and six were identified between unrelated neurons (0.0098%, 6 out of 612; *p*=0.43, Fisher’s exact test, Figure 4D).

However, when we considered only connections linking cells vertically, across layers, we found that connection probability was increased between related pairs compared to unrelated pairs (P=6.0% [28 out of 464] and P=2.7% [19 out of 711] for related and unrelated pairs, respectively; *p*=0.0056, Fisher’s exact test; Figure 4E), consistent with prior studies (Yu et al., 2009, Yu et al., 2012). A single vertical bidirectional connection was identified between related neurons (0.0047%, 1 out of 211 vertical pairs in which both directions of connectivity were tested), and none were identified between unrelated neurons (0.0%, 0 out of 333; *p*=0.39, Fisher’s exact test; Figure 4E). At the level of layer-specific connection types, we found that the connection probabilities from L4 to L5 (P=11.3% [7 out of 62] and P=2.3% [2 out of 89] for related and unrelated pairs, respectively; *p*=0.033, Fisher’s exact test; Figure 4F) and from L2/3 to L5 (P=7.9% [6 out of 76] and P=1.6% [2 out of 123], for related and unrelated pairs respectively; *p*=0.056) were increased between related compared to unrelated pairs.

To estimate the contribution of clonally related neurons as a fraction of the total input a cell receives, we used a simple quantitative model of connectivity based on our empirically measured connection probabilities and clone sizes. Briefly, we modeled the number of input connections to a particular cell from both related and unrelated cells in different layers of a cortical column as a binomial distribution with the connection probabilities set to the empirically measured connection probabilities (see Methods for additional assumptions). In particular, we wondered whether specific layer-defined connection types would show a substantial fraction of input from clonally related cells. Using our empirically measured connection probabilities and clone size (86 neurons on average), the model estimates that a substantial fraction of inputs from L4 to L2/3, L4 to L5, and L2/3 to L5 originates from cells with a common developmental lineage (7.2±2.6% of L4➔L2/3 connections, 19.2±12.2% of L4➔L5 connections, and 18.6±12.2% of L2/3➔L5 connections; estimate±SE, Figure 4G). These findings suggest that cell lineage may play an important role in shaping specific vertical, across-layer connections.

### Lateral, within-layer connections are not increased between excitatory neurons in radial clones

In contrast to vertical connections, we did not find evidence for an increase in the number of lateral connections within the same cortical layer between related neurons compared to unrelated neurons (P=5.7% [14 out of 248 potential connections] and P=8.2% [51 out of 626 potential connections] for related and unrelated pairs, respectively; *p*=0.25, Fisher’s exact test; Figures 5A–C). There was also no statistically significant difference in bidirectional lateral connections between related and unrelated pairs (0%, 0 out of 113 related lateral pairs in which both directions of connectivity were tested; 2.1%, 6 out of 284 unrelated lateral pairs in which both directions of connectivity were tested; *p*=0.19, Fisher’s exact test; Figure 5C). If anything, related neurons were more rarely connected to each other within L2/3 or within L5 (P=1.9% [2 out of 105 potential connections] and P=2.3% [1 out of 43 potential connections] for related pairs within L2/3 and L5, respectively) compared to unrelated pairs (P=5.9% [20 out of 342 potential connections] and P=10.3% [14 out of 136 potential connections] for unrelated pairs within L2/3 and L5, respectively) although the differences were not statistically significant (*p*=0.13 and *p*=0.12 for L2/3 and L5, respectively, Fisher’s exact test; Figure 5D). Our connectivity model estimates that only a small fraction of lateral, within-layer input to a cell comes from related neurons (1.5±1.1% of connections within L2/3, 4.3±1.5% of connections within L4, and 1.1±1.1% of connections within L5; estimate±SE; Figure 5E). Overall, these data suggest a revised model of circuit assembly among clonally related excitatory neurons in which related cells are preferentially connected vertically, across cortical layers, but not within a layer (Figure 5F).

**Figure 5.**
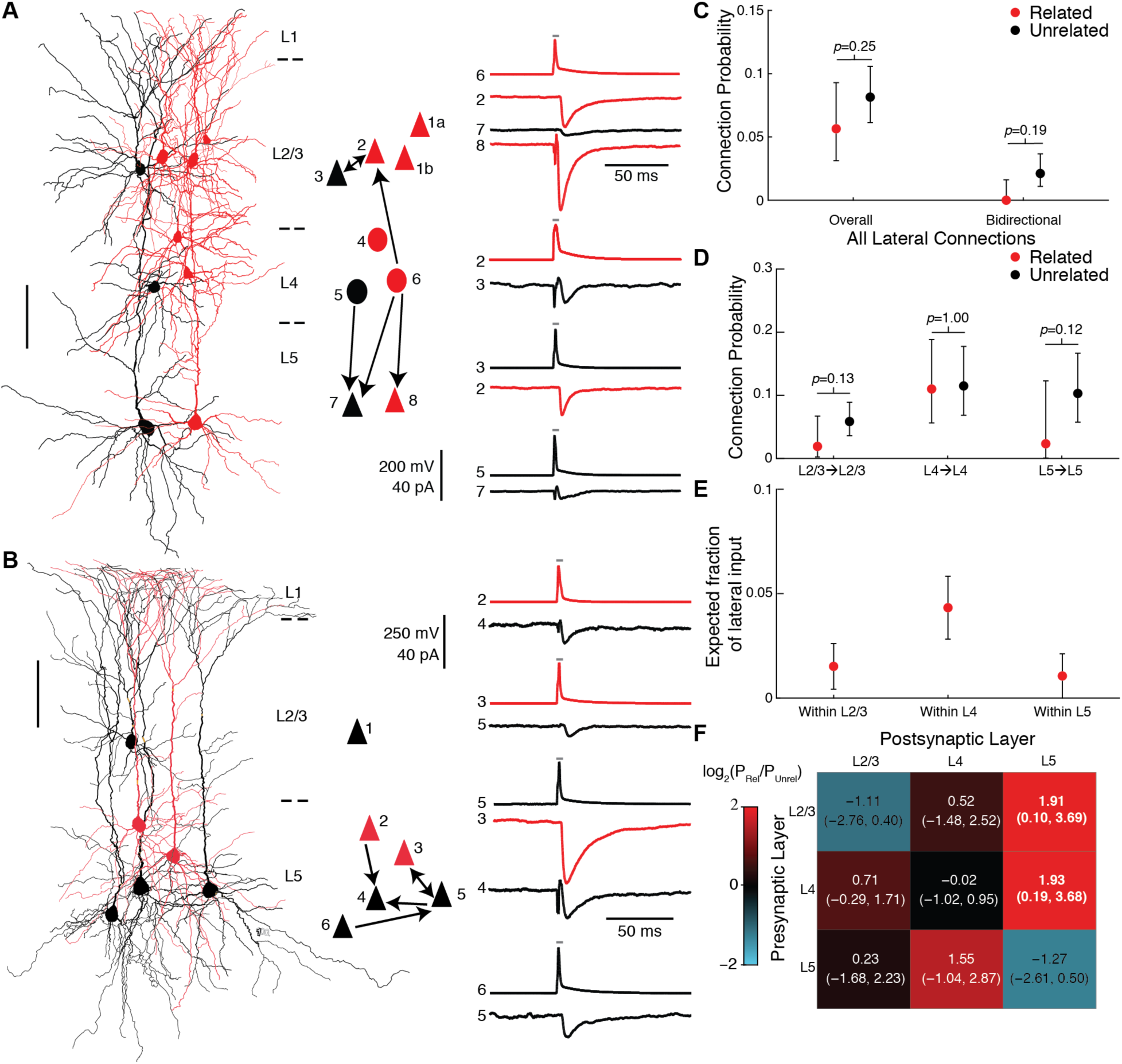
Lateral, within-layer connections are not increased between excitatory neurons in radial clones. **(A and B)** Example recording sessions testing within-layer connections among clonally related cells (red) and nearby, unrelated control cells (black) within L2/3 **(A)** and L5 **(B)**. Scale bars, 100 µm. Triangles, pyramidal neurons; ovals, L4 excitatory neurons. Presynaptic action potential (AP) and postsynaptic uEPSC traces for each connection are an average of at least 20 trials each. Grey bar indicates period of depolarizing current injection to presynaptic neuron. **(C)** Connection probabilities among related and unrelated neurons, pooling all lateral, within-layer connections tested (n=248 potential connections and 113 pairs with both directions tested for related neurons; n=626 potential connections and 284 pairs with both directions tested for unrelated neurons). **(D)** Connection probabilities among related and unrelated neurons, for each lateral connection type tested (n=105, 100, and 43 potential connections for related neurons and n=342, 148, and 136 potential connections for unrelated neurons within L2/3, L4, and L5, respectively). **(E)** Estimated fraction of lateral inputs to a single cell within L2/3, L4, or L5 that comes from clonally related neurons based on our empirically measured clone sizes and connection probabilities. **(F)** Heatmap of the log ratio of the connection probabilities for related and unrelated neurons, with additive smoothing (〈=1) by connection type tested. For **(C–D)**, error bars are 95% Clopper-Pearson confidence intervals and *p*-values are computed using Fisher’s exact test. For **(E)**, error bars are propagated standard error of the estimates (see Methods) For **(F)**, the 95% confidence interval, in parentheses below the value, is computed by resampling; significant values are highlighted in bold. See also Figures S7 and S8.

Since connection probability also depends on the distance between cells (Perin et al., 2011, Ko et al., 2011), and potentially on cortical area, we performed additional analyses in order to take these variables into account. First, for each pair of clonally related neurons we identified a set of matched control pairs with the same pre- and post-synaptic layers and with the same (up to 20 µm difference) tangential and vertical distances between the cells. We compared connectivity rates between related pairs and distance-matched control pairs using bootstrapping (see Methods). We found similar results as described above, with increased vertical connection probability between related neurons and no evidence for a difference in lateral connection probability (Figure S6). Second, we sorted the data into two groups according to the rostro-caudal position of each clone, which revealed similar changes in connectivity between related and unrelated neurons in both groups (Figure S7). Third, we built a generalized linear model of connection probability depending on the cell lineage (related or unrelated), connection type (vertical or lateral), Euclidean distance between cells and rostro-caudal position as predictors (see Methods and Table S3). The model revealed a significant interaction between cell lineage and connection type (*p*=0.035), further supporting that the effect of cell lineage on connectivity depends on the type of connection tested. The model also revealed a weak decrease in connection probability with Euclidean distance for lateral connections (*p*=0.056) and no evidence for any influence of rostro-caudal position (*p*>0.5 for the main effect and all interactions).

## Discussion

In summary, we show that radial clones in the mouse neocortex are composed of a diverse ensemble of excitatory neurons that are more likely to be synaptically connected by vertical connections across cortical layers, but not by lateral connections within a cortical layer. These findings carry important implications regarding the developmental mechanisms of circuit assembly in the neocortex and suggest that integration of vertical input from related neurons with lateral input from unrelated neurons may represent a fundamental principle of cortical information processing that is initially established by hardwired developmental programs.

### Cell type composition of radial clones

All of the radial clones labeled at E10.5 or earlier in our study spanned both superficial and deep layers of the cortex, consistent with prior studies labeling progenitors at this early developmental stage (Tan et al., 1998). Some groups have reported upper layer fate-restriction among a subset of radial glia when labeling progenitors at E10.5 (Franco et al., 2012, Franco and Muller, 2013, Gil-Sanz et al., 2015) using the *Cux2*-CreER driver line but not the Nestin-CreER line (Franco et al., 2012). It is possible that *Cux2*-CreER labels a small subset of radial glia that exhibit this neurogenesis pattern, or that the progenitor pool labeled using our *Nestin*-CreER driver is distinct from the *Cux2*-CreER-positive fraction. Given the low doses of tamoxifen administered in our study, we may be biased to labeling progenitors with the highest expression of *Nes* at E10.5, most likely ventricular radial glial cells (Hockfield and McKay, 1985), but potentially some symmetrically dividing neuroepithelial stem cells as well, which express low levels of Nestin in their pial end feet (Misson et al., 1988). It is also possible that the upper layer fate-restricted progenitors present at early time points, such as intermediate progenitors (Mihalas & Hevner, 2018), collectively generate all the different excitatory neuron subtypes within a given cortical region in parallel with the translaminar clones labeled in our study, which could explain why we saw no difference in the overall distribution of cell types between our labeled clones and randomly selected unlabeled excitatory neurons. Interestingly, the only transcriptomic subtype that appeared relatively underrepresented in our clones was a subset of L2/3 intratelencephalic neurons (L2/3 IT VISp Rrad; Figure S4B), but this difference was not statistically significant after correction for multiple comparisons. Further work is needed to determine whether this particular subtype of L2/3 neurons may arise primarily from *Cux2*-positive or other progenitors.

As this manuscript was being prepared, a preprint was published (Llorca et al., 2018) suggesting that individual translaminar clones are composed of restricted subtypes of excitatory neurons; in particular, they report approximately 10% of clones restricted to either the superficial or deep layers, and nearly a quarter of their translaminar clones were composed exclusively of corticocortical projection neurons. Two differences may explain this discrepancy. First, that study used a different Cre driver line (*Emx1*-CreERT2) which may target a different progenitor pool than the one described in our study. Second, their clones were labeled at a later developmental stage (E12.5), raising the possibility that their layer-restricted clones represent subclones that, if labeled earlier in neurogenesis, would give rise to clones spanning both superficial and deep layers and containing a mixture of cell types. Consistent with this interpretation, we saw that approximately one quarter of our clones were restricted to superficial layers when labeled at E11.5, but we never saw this upper-layer restriction when labeling at E9.5 or E10.5.

Recent single-cell RNA-sequencing studies have shown that excitatory neuron cell types are largely region-specific, at least between V1 and ALM (Tasic et al., 2018). Here we profiled the transcriptomes of cells from two primary sensory areas, V1 and S1, and found that the vast majority of cells from both regions map to V1-specific transcriptomic cell types rather than ALM-specific transcriptomic cell types, suggesting that cell types in different primary sensory areas are more similar to each other than they are to cell types in motor cortex. Interestingly, we still observed region-specific differences in gene expression between V1 and S1, suggesting that these two cortical areas are likely composed of distinct excitatory neuron types as well. While there was no evidence to suggest that the quality of our mapping to the V1/ALM reference dataset was worse for S1 cell than for V1 cells, if a reference atlas for S1 becomes available in the future it would be interesting to re-examine our data to better understand the developmental timeline of area-specific gene expression signatures in these two primary sensory areas.

### Connectivity matrix of radial clones

We find that excitatory neurons in radial clones are more likely to be synaptically connected vertically, across cortical layers, but not laterally, within the same cortical layer. While it has been previously reported that clonally related excitatory neurons are more likely to be synaptically connected (Yu et al., 2009, Yu et al., 2012, He et al., 2015), these prior studies did not tease out the effect of shared lineage on different types of layer-defined connections and, in particular, did not report any results for within-layer connections between clonally related neurons. Thus, our findings provide a higher resolution model of how cell lineage shapes developing cortical circuits.

A more recent study using chimeric mice with fluorescently labeled induced pluripotent stem cells (iPSCs) injected into blastocysts at E3.5 has examined lateral connections between related cells within L4 and found a transient increase in synaptic connectivity at P13-P16, which is followed by an increase in the fraction of connections that are reciprocal, rather than one-way, at P18-P20 (Tarusawa et al., 2016). We did not find any evidence for an increase in either overall connectivity or bidirectional connectivity in our data; however, it is possible that if the increase in connectivity in L4 is transient and only present from P13-P16 for overall connectivity and from P18-P20 for bidirectional connectivity that we may have missed it, as our data span the space between these time windows from P15-P20. Another possibility is that the iPSC-derived neurons may have altered synaptogenesis due to chromosomal instability and altered gene expression programs of iPSCs (Mayshar et al., 2010). Additional experiments to explore the possibility of a transient increase in lateral connections that include a direct comparison between iPSC-derived clones and clones labeled using other methods may ultimately resolve these questions.

Our finding that clonally related neurons are only rarely connected by lateral connections within L2/3 is particularly unexpected given prior studies showing more similar feature selectivity between clonally related neurons in this layer (Li et al., 2012), even in very large clones (Ohtsuki et al., 2012). Several studies have now shown that, within L2/3, excitatory neurons that have similar orientation tuning are more likely to be synaptically connected and have stronger synapses compared to cells with dissimilar tuning preferences (Ko et al., 2011, Cossell et al., 2015). Thus, the expectation and proposed model (Li et al., 2018) would be that increased connections between clonally related neurons within L2/3 underlies their similarity in tuning. However, we found no evidence for an increase in lateral connections between related cells in L2/3, suggesting that they may inherit similar feature selectivity either by receiving common feedforward inputs from L4 or by modulation from long-range feedback connections. Moreover, these results suggest a novel functional role for lateral connections within L2/3 in permitting synapse formation between cells in unrelated clonal units, in addition to linking cells with similar tuning preferences.

Our results highlight L5 as a potential hub within radial clones, with the most striking increases in connectivity seen in the projections from superficial layers to L5. Neurons in L5 serve as a major output of the cortex with important roles in integrating feedback from higher cortical areas and in top-down modulation by brain states (Kim et al., 2015) and altered gene expression in deep layer neurons during midfetal development has been recently implicated in neuropsychiatric disorders such as autism (Willsey et al., 2013). We propose that integration of translaminar input from clonally related neurons with intralaminar input from unrelated neurons in L5 may represent an organizing principle for lineage-dependent circuit assembly. While L5 has traditionally been less amenable to in vivo functional studies, recent advances in calcium imaging such as three-photon microscopy and genetically encoded calcium indicators (Ouzounov et al., 2017) may enable functional analysis of cortical computation in both superficial and deep layers of radial clones. Future studies aimed at dissecting the functional role of lineage-driven synaptic connectivity across the cortical column may provide mechanistic insight into abnormal circuit function in neuropsychiatric disease.

The mechanism by which radial clones of excitatory neurons form specific connections is thought to involve gap junction coupling during migration along the radial glial fiber (Yu et al., 2012). A recent study has further shown that it is the coupling between clonally related neurons, and not between the postmitotic neurons and their radial glia or progenitors, that promotes specific synapse formation between radially aligned sister neurons (He et al., 2015). This coupling requires the inside-out migration of related neurons along a similar path and is abolished by removal of REELIN or its downstream effector DAB1 which disrupt inside-out migration, or by increased levels of EFNA/EPHA-mediated signaling which leads to increased lateral displacement of clonally related neurons as they traverse the intermediate zone prior to reaching the cortical plate (Torii et al., 2009, He et al., 2015). Interestingly, early studies suggested that migration along multiple radial glial fibers may be common within radial clones (Walsh and Cepko, 1988) and underlie the substantial tangential dispersion seen within radial clones. Our finding that connections from L2/3 to L5 and from L4 to L5 are specifically enhanced between clonally related neurons could be consistent with a mechanism that requires inside-out migration along a radial glial fiber and further suggests that as migrating neurons travel to reach the superficial layers, their axons may “stick” to the maturing apical dendrites of clonally related deep layer neurons they are passing. However, we did not see any difference in connectivity between related neurons based on their tangential displacement (data not shown), as might be expected if radial migration along the same glial fiber is necessary for formation of specific synapses. The possibility that clonally related neurons may either establish specific vertical connections regardless of which radial glial fiber they follow, for example by expression of specific cell adhesion molecules during migration (Tarusawa et al., 2016), or undergo significant tangential migration *after* passing deep layer neurons may warrant further investigation.

It is nearly impossible to prove that an effect is absent, and with additional sampling of lateral connections, particularly in L4 as discussed above, it is possible that a difference in connectivity may emerge. However, our data suggest that any effect of lineage on lateral connectivity must be very small and, based on the trend seen in our data within L2/3 and within L5, may actually be in the opposite direction with fewer connections between clonally related neurons than between unrelated pairs. One possible explanation for this is that any two clonal related neurons in the same layer could be generated by a symmetrically dividing intermediate progenitor cell, but not radial glia, and it is theoretically possible that progenitor cell type of origin could influence development of local cortical microcircuit. Another possibility is that any two neurons in the same layer were generated by two different radial glia, which shared a common symmetrically dividing radial glia ancestor labelled at E10.5. Future studies utilizing temporally resolved lineage tracing methods (McKenna et al., 2016) could provide further insights into how the degree of relatedness impacts intra-clonal connectivity.

Similarly, since we focused on V1 and S1, both primary sensory areas, it is possible that a different pattern of connectivity among clonally related neurons is present in other cortical areas such as primary motor cortex. We also tested only on local connections that can be tested in an acute slice preparation. Given that transcriptomic cell type correlates with the long-range projection pattern of excitatory neurons (Tasic et al., 2018), our finding that individual clones contain multiple diverse transcriptomic types suggests that neurons within radial clones might also project to diverse targets. Additional experiments using different methods for lineage tracing and connectivity profiling, focusing on different brain regions and using adult animals, will be necessary to determine the generalizability of the connectivity pattern we describe here and delineate the long-range inputs and outputs of individual radial clones. However, our data suggest that the integration of feedforward, intra-columnar input with lateral, inter-columnar information may represent a developmentally programmed connectivity motif for the assembly of neocortical circuits.

## Acknowledgements

We thank members of the Tolias and Sandberg labs including Alexander Ecker, Jacob Reimer, Dimitri Yatsenko, Shan Shen, Emmanouil Froudarakis, Amy Morgan, Camila Lopez, Megan Rech, Shannon McDonnell, and Leonard Hartmanis for helpful discussions and technical support, and Tomasz Nowakowski for critical reading of the manuscript. This project was supported by the Optical Imaging and Vital Microscopy core at Baylor College of Medicine; grants R01MH103108, R01DA028525, DP1EY023176, P30EY002520, T32EY07001, and DP1OD008301 from the National Institutes of Health (NIH) to A.S.T.; grants from the Swedish Research Council and the Vallee Foundation to R. S.; grants from the Deutsche Forschungsgemeinschaft (DFG, EXC 2064, BE5601/4-1) and the German Federal Ministry of Education and Research (BMBF; FKZ 01GQ1601) to P.B.; the McKnight Scholar Award to A.S.T.; and the Arnold and Mabel Beckman Foundation Young Investigator Award to A.S.T. X. J. was supported by BCM Faculty Start-up Fund. C.R.C was supported by NIH grants F30MH095440, T32GM007330 and T32EB006350. P.G.F. was supported by NIH grant F30MH112312. C.R.C and P.G.F. were both supported by the Baylor Research Advocates for Student Scientists (BRASS) foundation.

This project was also supported by the Intelligence Advanced Research Projects Activity (IARPA) via Department of Interior/Interior Business Center (DoI/IBC) contract number D16PC00003. The U.S. Government is authorized to reproduce and distribute reprints for Governmental purposes notwithstanding any copyright annotation thereon. Disclaimer: The views and conclusions contained herein are those of the authors and should not be interpreted as necessarily representing the official policies or endorsements, either expressed or implied, of IARPA, DoI/IBC, or the U.S. Government.

This project was also supported by the National Institute of Mental Health and the National Institute of Neurological Disorders and Stroke under Award Number U19MH114830. The content is solely the responsibility of the authors and does not necessarily represent the official views of the National Institutes of Health.

## Author Contributions

C.R.C. and P.G.F. generated mice with labeled clones. X.J., F.S. and S.L. performed electrophysiological recordings for Patch-seq experiments. C.R.C. and F.S. amplified full-length cDNA from patched cells. C.R.C. generated sequencing libraries and performed pre-processing of the sequencing data with assistance from P.J. X.J. and F.S. performed the multi-patching experiments. C.R.C. performed quantitative analysis of clones and analyzed Patch-seq data with input from D.K. D.K. mapped Patch-seq cells to the reference dataset and performed joint t-SNE projections. C.R.C., X.J., and F.S. analyzed the connectivity data with input from D.K. and R.J.C. F.H.S. implemented the connectivity model. R.S. supervised the library preparation, sequencing, and pre-processing of sequencing data. P.B. supervised all data analysis. X.J. supervised the multi-patching and Patch-seq experiments. A.S.T. supervised all experiments and analyses. C.R.C. drafted the manuscript with input from all co-authors.

## Methods

### Animals

All experiments were carried out in accordance with, and with approval from, the Institutional Animal Care and Use Committee (IACUC) at Baylor College of Medicine (BCM). The Nestin-CreER line was obtained from M. Maletic-Savatic (BCM) and maintained in A. Tolias’ laboratory by crossing heterozygous males with wild type C57Bl/6J females. Each generation, potential stud males were crossed with a reporter line to confirm the lack of transgene expression in the absence of tamoxifen administration, and only those males showing minimal to no “leaky” recombination in the P1 offspring of this test cross were used as breeders for maintaining the Cre line. The Nestin-CreER line will be cryopreserved at BCM for potential future use. The reporter line ROSA26-CAG-LSL-tdTomato-WPRE (Ai9) was acquired from the Jackson Laboratory (JAX Stock #007909). The outbred CD1 line was obtained from the Center for Comparative Medicine at BCM. Six mice were used for quantification of clone size and width (3 males and 3 females), 9 mice (all males) were used for Patch-seq experiments, and 43 mice (26 males, 7 females, 10 uncertain) were used for electrophysiology experiments. For clone quantification, animals were sacrificed at postnatal day 10 (P10) and for Patch-seq and multi-patching experiments animals were sacrificed at P15-P20. All animals were on the C57Bl/6J or mixed C57Bl/6J; CD1 genetic background and were group housed with their littermates and foster mothers (both CD1 and C57Bl/6J foster mothers were used) on a 12-hour light-dark cycle.

### Lineage tracing

We used a tamoxifen-inducible Cre-lox transgenic approach for lineage tracing similar to previous studies (Gao et al., 2014). Two breeding strategies were used: The majority of experimental animals were generated by crossing *Nestin*-CreER heterozygous males with Ai9 homozygous females. A minority of experimental animals were generated by crossing double homozygous Cre; Ai9 males (C57/Bl6J) with wild type CD1 females. The latter breeding strategy negated the need for genotyping of the pups (all would be double heterozygotes) and substantially increased litter size.

Tamoxifen and progesterone were dissolved together in corn oil and administered to pregnant dams at E9.5, E10.5 or E11.5 at a dose of 40–50 and 20–25 mg/kg, respectively, by orogastric gavage. To help prevent tamoxifen-induced pregnancy loss (Milligan & Finn, 1997), pregnant mice also received 2 mg of progesterone dissolved in corn oil subcutaneously twice a day, starting the day after tamoxifen treatment and continuing until the pups were delivered by Caesarian section on E19.5 (as described in Nagy et al., 2006). The pups were raised by a foster mother and standard genotyping protocols were used to identify double heterozygous animals carrying both the Cre and reporter alleles, if needed depending on the breeding strategy.

### Transcardial perfusion and histology for clonal analysis

Animals were deeply anesthetized with isoflurane and transcardially perfused with 0.1M phosphate buffered saline (PBS) followed by 4% paraformaldehyde in PBS at postnatal day 10 (P10). Fixed brains were coronally sectioned at 100 µm on a vibratome (Leica VT1000S) and stained with DAPI (0.25 µg/mL) for 10–15 min before mounting on charged glass slides with anti-fade mounting solution (1 mg/ml ρ-phenylenediamine in 90% glycerol, 10% PBS, pH ~8.0). Confocal image stacks were taken on either a Zeiss LSM 510 Meta or a Zeiss LSM 780 confocal microscope.

### Acute brain slice preparation

Acute brain slices were prepared as previously described (Jiang et al., 2015). In brief, animals (P15–P20) were deeply anesthetized with 3% isoflurane and decapitated. The brain was quickly removed and placed into cold (0–4 °C) oxygenated physiological solution containing (in mM): 125 NaCl, 2.5 KCl, 1.25 NaH2PO4, 25 NaHCO3, 1 MgCl2, 25 dextrose, and 2 CaCl2, pH 7.4. Parasagittal slices 300 µm thick were cut from the tissue blocks using a microslicer (Leica VT 1200). The slices were kept at 37.0±0.5°C in oxygenated physiological solution for ~0.5–1 h before recordings. During recordings, the slices were submerged in a chamber and stabilized with a fine nylon net attached to a platinum ring. The recording chamber was perfused with oxygenated physiological solution. The half-time for the bath solution exchange was 1–2 min, and the temperature of the bath solution was maintained at 34.0±0.5°C. All antagonists were bath applied.

### Patch-seq sample collection

To obtain transcriptome data from individual neurons within radial clones, we used our recently described Patch-seq method (Cadwell et al., 2016, Cadwell et al., 2017). Briefly, the following modifications were made to the standard whole-cell patch-clamp workflow to improve RNA yield from patched cells. Glass capillaries were autoclaved prior to pulling patch pipettes, all work surfaces and micromanipulator pieces were thoroughly cleaned with DNA-OFF and RNase Zap, and all solutions that would come into contact with RNA were prepared using strict RNAse-free precautions. Recording pipettes of 2–4 ΜΩ were filled with a small volume (approximately 0.3µl) of intracellular solution containing: 123 mM potassium gluconate, 12 mM KCl, 10 mM HEPES, 0.2 mM EGTA, 4 mM MgATP, 0.3 mM NaGTP, 10 mM sodium phosphocreatine, 20 µg/ml glycogen, 13 mM biocytin, and 1 U/µl recombinant RNase inhibitor, pH ~7.25. RNA was collected at the end of whole-cell recordings by applying light suction while observing the cell under differential interference contrast (DIC). If any extracellular contents were observed to enter the pipette under DIC, the sample was discarded. Otherwise, the contents of the pipette were ejected into and RNase-free PCR tube containing 4 µl of lysis buffer consisting of: 0.1% Triton X-100, 5 mM (each) dNTPs, 2.5 µM Oligo-dT30VN, 1 U/µl RNase inhibitor, and 1×10^−5^ dilution of ERCC RNA Spike-In Mix.

### cDNA synthesis, library preparation and sequencing

Single cell RNA was converted to cDNA following the Smart-seq2 protocol (Picelli et al., 2014a, Cadwell et al., 2017). Samples were denatured at 72°Cfor 3 min and then 5.70 µl of RT mix was added to each sample, for final concentrations of: 1× Superscript II first strand buffer, 1M Betaine, 10U/µl SSIIRT, 5 mM DTT, 1 U/µl RNase inhibitor, 1 µM LNA-TSO, and 6 mM MgCl2. The RT reaction was run at 42°Cfor 90 min followed by ten cycles of 50°Cfor 2 min, 42°Cfor 2 min, and the enzyme was inactivated by holding at 70°C for 15 min.

The full-length cDNA was amplified by adding 15 µl of PCR mix to each sample, consisting of 1× KAPA HiFi HotStart Ready Mix and 0.1 µM IS PCR primers, and running the following PCR program: 98°Cfor 3 min; 18 cycles of 98°Cfor 20 s 67°Cfor 15 s 72°Cfor 6 min; and 72°Cfor 5 min. The PCR product was purified using Axygen AxyPrep mag PCR beads according to the manufacturer’s instructions but using a bead:sample ratio of 0.7:1 (17.5 µl of beads: 25 µl sample).

To construct the final sequencing libraries, we diluted each sample to a concentration of 50 pg/µl and added 4 µl of tagmentation mix to 300 pg (6 µl) of full-length cDNA for a final concentration of: 1×tagmentation buffer (1mM TAPS-NaOH, 5 mM MgCl2), 10% (wt/vol) PEG-8000, and 1.25 µM in-house produced Tn5 transposase (Picelli et al., 2014b, Cadwell et al., 2017). The tagmentation reaction was run in a thermal cycler at 55°Cfor 8 min and the Tn5 transposase was stripped by adding 2.5 µl of 0.2% (wt/vol) SDS to each sample by incubating at room temperature for 5 min.

Amplification of the adapter ligated fragments was performed by adding 2.5 µl each of index 1 (N7XX) and index 2 (N5XX) primers, diluted 1:4, from the Nextera XT index kit with a unique combination of indices for each sample, as well as 5 µl of 5× KAPA HiFi Buffer, 0.75 µl of KAPA dNTP mix (10 nM each), 1.25 µl of nuclease-free water, and 0.5 µl of KAPA enzyme (1U/µl) for a total volume of 25 µl. The enrichment PCR was run according to the following program: 72°Cfor 3 min, 95°Cfor 30 s, 12 cycles of 95°Cfor 10 s, 55°Cfor 30 s, and 72°Cfor 30 s, and 72°Cfor 5 min. After enrichment PCR, 2.5 µl of each library was pooled into a single 1.5 mL tube and purified using the Axygen AxyPrep mag PCR beads with a bead:sample ratio of 1:1. The pooled library was diluted to 3 nM and sequenced on a single lane of an Illumina HiSeq 3000 with single-end (50 bp) reads.

### Multi-cell recordings

Simultaneous whole-cell in vitro recordings were obtained from cortical neurons as previously described (Jiang et al., 2015). Briefly, patch recording pipettes (5–7 MΩ) were filled with intracellular solution containing 120 mM potassium gluconate, 10 mM HEPES, 4 mM KCl, 4 mM MgATP, 0.3 mM Na3GTP, 10 mM sodium phosphocreatine, Alexa-488 (10 µM) and 0.5% biocytin (pH 7.25). Whole-cell recordings were made from up to eight neurons simultaneously using two Quadro EPC 10 amplifiers (HEKA Electronic, Lambrecht, Germany). A built-in LIH 8+8 interface board (HEKA) was used to achieve simultaneous A/D and D/A conversion of current, voltage, command and triggering signal for up to eight amplifiers. Micromanipulators (Luigs & Neumann) were mounted on a ring specifically designed for multi-patching. PatchMaster software and custom-written Matlab-based programs were used to operate the Quadro EPC 10 amplifiers and perform online and offline analysis of the data. In order to reveal passive membrane properties and firing patterns of each recorded neurons, neurons were stimulated with 600 ms long current pulses starting from -100 / -200 pA with 20 pA steps.

Recordings were made in cortical layers 2/3, 4 and 5, targeting fluorescently labeled (red) cells as well as nearby unlabeled neurons that had clear pyramidal somata and apical dendrites, with the exception of neurons in L4. We visually confirmed successful targeting of tdTomato-expressing neurons based on the spatial overlap of green (due to Alexa-488 in patch pipette) and red fluorescence (see Figure 4B). Unitary excitatory postsynaptic currents (uEPSCs) were evoked by current injection into the presynaptic neurons at 2–3 nA for 2 ms while clamping or holding the membrane potential of the postsynaptic cells at −70mV. Each neuron was assigned to a laminar position using layer boundaries visible in the high-contrast micrographs obtained during electrophysiological experiments, and confirmed post-hoc using the recovered morphology. Latency was defined as the time from the peak of the presynaptic action potential (AP) to 5% of the maximum amplitude of the uEPSC. Amplitude was defined as the maximum amplitude of the uEPSC from baseline. Latency and amplitude are reported as mean±SD across all connections analyzed.

### Morphological reconstruction after whole-cell recordings

Light microscopic examination of the morphology of each neuron was carried out following previously described protocols (Jiang et al., 2015, Cadwell et al., 2017). In brief, after in vitro recordings, the slices were fixed by immersion in 2.5% glutaraldehyde/4% paraformaldehyde in 0.1 M phosphate buffer at 4°C for at least 48 h, and then processed with an avidin-biotin-peroxidase method to reveal cell morphology. The morphologically recovered cells were examined using a 100× oil-immersion objective lens and camera lucida system (Neurolucida, MBF Bioscience). In addition, the 3D coordinates of the cells were measured and the distance between each pair of simultaneously recorded neurons was computed, including Euclidean distance, tangential distance (parallel to pial surface) and vertical distance (perpendicular to pial surface).

### Quantification of clone size and width

To quantify the width and number of neurons per clone, six near-complete brain sets were analyzed using custom-written Matlab software and manual cell segmentation (two brains each treated at E9.5, E10.5 or E11.5, continuous sections spanning 3–4 mm along the rostrocaudal axis) as follows: a two-dimensional maximum projection of each slice was divided into small sections and presented one at a time to a blinded observer for manual identification of neurons throughout the cortex. Glia were excluded based on morphology. Clones were manually reconstructed across slices by aligning fiducial anatomic landmarks such as the longitudinal fissure. The number of neurons within each clone was calculated by adding together all of the neurons within the clone across all contiguous slices where the clone was identifiable. On each slice, the widest part of the clone was measured, and the overall width for each clone was computed as the median of the measured width of the clone across all slices. Clone width and number of neurons per clone are reported as the median and interquartile range (IQR) in the text and all of the individual data points are shown in Figures 1D-1E. The number of clones and animals for each treatment condition are reported in the figure legend. The Wilcoxon rank sum test was used to compare E9.5 to E10.5 and E10.5 to E11.5 and those *p*-values are also shown in Figures 1D-1E.

Clones were also classified as complete or incomplete after reconstruction by a blinded observer based on whether they spanned all cortical layers, including L5 and L6. All clones that were considered “incomplete” are shown in either Figure 1C or Figure S1A. The fraction of all clones that were considered incomplete for each treatment condition is reported in Figure 1F as well as the 95% Clopper-Pearson confidence intervals for each ratio. The number of clones and animals for each treatment condition are reported in the figure legend. Fisher’s exact test was used to compare E9.5 to E10.5 and E10.5 to E11.5 and those *p*-values are also shown in Figure 1F.

### Single-cell RNA-sequencing analysis

#### Quality control and data pre-processing

A total of 278 neurons from 16 radial clones were aspirated for single-cell RNA-sequencing experiments. The quality of the full-length cDNA for each sample was analyzed by running on and Agilent bioanalyzer with a High Sensitivity DNA chip. Samples containing less than ~1 ng total cDNA (less than ~67pg/µl) or with an average size less than 1,500 bp when integrating over the range from 300 to 9,000 bp were not sequenced (~21%, 58/278 neurons, leaving 220 samples).

The final pooled sequencing library was also analyzed on an Agilent Bioanalyzer to confirm that the average library size was less than ~500 bp and there were minimal primer dimers. Reads were aligned to the mouse genome (mm10 assembly) using STAR (v2.4.2a) with default settings. Only read counts were used for the data analysis presented here. Eleven cells were excluded after sequencing due to poor quality sequencing results (~5%, 11/220 neurons, leaving 209 samples; poor quality was defined as greater than three median absolute differences below the median for either total number of reads or total number of genes detected; Figures S3A-B). Three additional neurons (~1.4%, 3/209) were excluded from further analysis because they had fast-spiking or regular-spiking firing patterns consistent with inhibitory interneurons, leaving 206 samples for all subsequent analyses.

Genes with less than one read per cells on average (Figure S3C) were removed (*n*=12,841 genes remaining) and the count data were normalized using the scran package in R Bioconductor (Lun et al., 2016). Quality control plots (Figures 2C-E, S3 and S4) were performed using scran as described in Lun et al., 2016. Across genes, there was a strong correlation between the average count per cell and the number of cells expressing each gene (Figure S3D) and alternatively filtering genes based on the number of cells expressing each gene had no significant effect on our results (data not shown). The normalized read counts were used for all subsequent analyses.

#### Dimensionality reduction within our dataset

To reduce the dimensionality for visualizing gene expression within our own dataset, we used the R Bioconductor implementation of t-distributed Stochastic Neighbor Embedding (t-SNE, runTSNE function of the scran package) with the random seed set to 30 for reproducibility. As input, we used the normalized and log2-transformed counts (“logcounts”, Table S1) of the top highly variable genes selected with a false discovery rate set to 0.05 (computed using the correlatePairs function with per.gene=TRUE) among the cells being plotted (n=91 genes for Figures 2F and S3C, n=43 genes for Figure 2G and n=41 genes for Figure 2H). The parameter for perplexity was set to 30 when analyzing all cells (Figures 2F and S3C), to 10 when analyzing only L2/3 cells (Figure 2G), and to 15 when analyzing only L5 cells (Figure 2H). Very similar two-dimensional projections were generated when different parameters or number of genes were used.

#### Generalized linear models (GLMs) to predict layer and region

We used the cv.glmnet function in R Bioconductor to train a GLM to predict either layer (Figure 2F right panel) or cortical region (Figures 2G and 2H, right panels) as follows:

~~~
cvfit<-cv.glmnet(logcounts,factor,family="multinomial",parallel=TRUE,type.mea sure="class",nfolds=20)
~~~

The model performance was estimated from the lowest prediction error across all lambda values as follows:

~~~
perc_correct <-1-cvfit$cvm[which(cvfit$lambda==cvfit$lambda.min)]
~~~

To generate a null distribution for each model, we randomly shuffled either the layer position (for Figure 2F) or cortical region (for Figures 2G and 2H) by resampling without replacement 1000 times. For each iteration, the model performance was evaluated as described above. The *p*-values are computed as the fraction of resamples with model performance (percent correct) greater than or equal to the unshuffled model performance. The values in the rightmost panels of Figures 2F-H are the unshuffled model performance (in black) and the mean and 95% coverage interval of the resampled model performances (in grey).

#### Mapping to the reference dataset using t-SNE

Using the count matrix of Tasic et al. 2018 (*n*=23,822, *d*=45,768), we selected 3000 “most variable” genes as described in Kobak & Berens, 2018. Briefly, we found genes that had, at the same time, high non-zero expression and high probability of near-zero expression. In particular, we excluded all genes that had counts of at least 32 in fewer than 10 cells. For each remaining gene, we computed the mean log2 count across all counts that were larger than 32 (non-zero expression, *µ*) and the fraction of counts that were smaller than 32 (probability of near-zero expression, *τ*). Across genes, there was a clear inverse relationship between *µ* and *τ,* that roughly followed an exponential law *τ ≈* exp*(-µ+a)* for some horizontal offset *a.* Using a binary search, we found a value *b* of this offset that yielded 3000 genes with *τ >* exp*(-µ+b) + 0.02.* These 3000 genes were selected as input for dimensionality reduction.

The t-SNE visualization of the Tasic et al. 2018 dataset shown in Figures 3A S3A, and S4 was generated as described in our previous work (Kobak & Berens, 2018). It was computed there using PCA initialization and perplexity combination of 50 and 500, following preprocessing steps of library size normalization (by converting counts to counts per million), feature selection (using the 3000 most variable genes), log2(x+1) transformation, and reducing the dimensionality to 50 using PCA. The resulting t-SNE coordinates for all Tasic et al. cells are given in Table S2.

Out of 3000 most variables genes selected in the Tasic et al. data set, 1181 genes were present among the 12,841 that we selected in our data set. We used this set of 1181 genes for the mapping of our cells to the reference data. For each of the *n*=206 Patch-seq cells in our dataset, we computed its Pearson correlation with each of the 23,822 reference cells across the 1181 genes, after all counts were log2(x+1) transformed. We identified the 25 reference cells with the maximal correlation (25 “nearest neighbors” of our cell) and positioned our cell at the median t-SNE location of those 25 reference cells (Kobak & Berens, 2018).

We performed bootstrapping over genes to estimate the uncertainty of this mapping (Kobak & Berens, 2018). Specifically, we selected a bootstrap sample of 1181 genes and repeated the mapping as described above. This was repeated 100 times, to obtain 100 bootstrap positions of each cell. We computed the Euclidean distance between the original mapping position and each of the bootstrap positions, and took the 80th percentile of the resulting distribution as a measure of mapping precision. If all bootstrap positions are close each to each other, the 80th percentile distance will be small (high precision). If they are far from each other, it will be large (low precision). This measure was used in Figures 3A and S3A (cells with the 80th percentile above 10 were plotted as small dots, cells with the 80th percentile greater than 5 but less than or equal to 10 were plotted as intermediate size dots, and cells with the 80th percentile less than or equal to 5 were plotted as large dots) and also to compare the quality of the mapping between V1 and S1 cells (see Results).

#### Mapping to the reference clusters

To assign each of our Patch-seq cells to one of the reference clusters, we log-transformed all counts from Tasic et al., 2018 with log2(x+1) transformation and averaged the log-transformed counts across all cells in each of the 133 clusters to obtain reference transcriptomic profiles of each cluster, using the same 1181 genes as above (133×1181 matrix). We applied the same log2(x+1) transformation to the read counts of our Patch-seq cells, and for each cell computed Pearson correlation across the 1181 genes with all 133 Tasic et al. 2018 clusters. Each cell was assigned to the cluster to which it had the highest correlation (nearest centroid classifier).

#### Probability of related and unrelated neurons mapping to the same clusters

To compute the probability related and unrelated pairs of neurons mapping to the same clusters (Figures 3C 3D, S3C, and S3D), we computed the number of pairs mapping to the same cluster as a fraction of all of the pairs analyzed. For Figures 3C and 3D, we first grouped the 133 transcriptomic clusters into ten broad classes, as labeled in Figure 3A. In the supplementary Figures S3C and S3D we kept all 133 original clusters. In Figures 3C and S3C, we included all pairs of neurons, and in Figures 3D and S3D we included only pairs of neurons that were positioned within the same cortical layer. The values shown are the overall fraction of pairs mapping to the same cluster or broad class, and the 95% Clopper-Pearson confidence intervals. The *p*-values are computed using the Chi-squared test.

### Comparison of connection probability between related and unrelated neurons

#### Comparison using raw data

Related pairs were defined as pairs in which both the pre- and post-synaptic neuron were tdTomato-positive excitatory neurons organized in a well-isolated radial unit of labeled cells (>300µm separation from other labeled clones). Control pairs were defined as pairs of nearby excitatory neurons in which one cell was tdTomato-positive (either the pre- or post-synaptic cell, but not both) and one was tdTomato-negative. The connection probability was determined as the total number of connections divided by the total number of connections tested within each category (all connections tested, only vertical connections, only lateral connections, and each layer-defined connection type). The values shown in Figures 4D-F and 5C-D are the connection probability and 95% Clopper-Pearson confidence intervals. The number of connections tested for each category is reported in the figure legends. Fisher’s exact test was used to compare the connection probabilities between related and unrelated cells and those *p*-values are shown in Figures 4D-F and 5C-D.

#### Comparison to distance-matched controls

Related pairs were defined as above. In contrast to the above comparison, control pairs were defined as pairs of excitatory neurons in which one or both cells were tdTomato-negative, to increase the number of available controls for distance-matching (a caveat is that two tdTomato-negative cells can in principle belong to another clone, but we consider this to be unlikely). For each related pair, we defined a set of “matched” control pairs for which the pre- and post-synaptic neurons were located in the same cortical layers as the pre- and post-synaptic neurons of the related pair, and for which both the tangential and vertical distances between the control cells were within 20 µm of the analogous distances between the two related cells. Related pairs that did not have any matching control pairs fitting these criteria were excluded from further analysis.

To compare connectivity between related and distance-matched control pairs, we used bootstrapping over related pairs. Specifically, on each of the 1000 iterations, we drew a bootstrap sample (resample with replacement) from the set of related pairs, and selected one matched control pair for each related pair. Values in Figure S6 are the mean and 95% coverage intervals across resamples of related and matched control connection probabilities. For each resample, we also computed the difference between the related and matched control connection probabilities. We “inverted” the bootstrap confidence interval for this difference to estimate the *p*-value. Specifically, the mean difference in connection probability was first subtracted from all bootstrapped differences, and the *p*-value was estimated as the fraction of resampled differences with absolute value greater than or equal to the original mean difference (two-tailed test). The *p*-values are shown in Figure S6.

#### Comparison at different rostrocaudal positions

To determine whether the effect of cell lineage varied according to rostrocaudal position, clones were sorted into two groups based on their rostrocaudal position (“rostral” includes clones within S1 proper but also other rostral cortical areas, and similarly for “caudal” clones and V1). The values shown in Figure S7 are the connection probability and 95% Clopper-Pearson confidence intervals for each group. The number of connections tested in each category is reported in the figure legend. Fisher’s exact test was used to compare the connection probabilities between related and unrelated cells in each group and those *p*-values are shown in Figure S7.

#### Generalized linear model of connection probability

We also built a generalized linear model (GLM) to explain connection probability (*P*) as a function of connection class, lineage relationship, Euclidean distance between the cells, and rostrocaudal position (a numeric value ranging from 1 to 5 with 1 being most rostral and 5 being most caudal). We fit a binomial GLM (using glmfit function in Matlab) containing the relevant linear terms and all possible pairwise interactions:

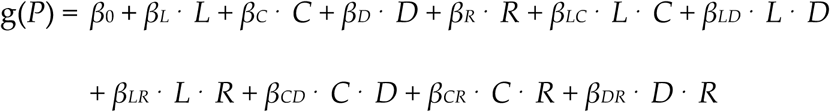

where *β*0 is a constant term, *L* is a binary variable representing the lineage relationship (1 for related and 0 for unrelated), *C* is a binary variable representing the connection type (1 for vertical and 0 for lateral), *D* is the Euclidean distance between the cells in microns, *R* is a numeric variable representing the rostrocaudal position of the clone with integer values from 1 (most rostral) to 5 (most caudal), and *βi* are the corresponding coefficients. The presence of a connection was modeled as Bernoulli distributed with probability *P*, using the logit link function, g(*P*) = ln(*P*/(1-*P*)). The estimated coefficients and *p*-values of each term are reported in Table S3.

### Simple connectivity model to estimate expected input from related cells

For a particular postsynaptic cell in layer *j* ∈ {L2/3, L4, L5}, we modeled the number of input connections from cells in a particular layer *i* and a particular lineage relation *l* ∈ {related, unrelated} as a binomial distribution B(*n_il_*,*p_il_*). The probabilities *p_ijl_* were set to the measured connection probabilities. The pool sizes *n_il_* were set to the product *n_il_* = *n_i_q_l_* of the number of cells *n_i_* residing in the particular input layer and the fraction *q_l_* of cells with that particular lineage. To compute *n_i_*, we assumed that a cortical slab of 1mm^2^ contains about 100,000 neurons and that 80% of those are excitatory neurons. We further assumed that a particular cell only connects to other cells within a tangential radius of *r*, which we set to half the 99% quantile of pairwise distances measured in our dataset (*r* = 0.087 mm). The resulting cylinder of cortex contained π*r*^2^ × 80,000 ≈ 1,908 excitatory neurons. We assumed that 35% of these cells reside in L2/3, 15% in L4, 25% in L5, and 25% in L6. The fraction *q*_c_ of related cells in that cylinder was computed as the ratio 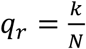 of the median clone size (*k* = 86) and the number of cells in the cortical cylinder (*N* = 1908). All model computations were performed using Python.

We computed the expected fraction of related cells in the input connections to a particular cell (Figures 4G and 5E) as

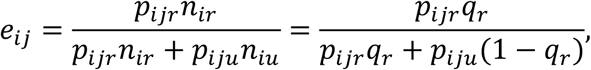

where subscript *r* refers to related neurons and *u* to unrelated neurons. Note that *q*_u_ = 1 − *q*_r_. We propagated the standard error from *p*_ijl_ to *e*_ij_ using a first order Taylor approximation: In general, the propagated variance of a function *f(X,Y)* of two random variables is given by

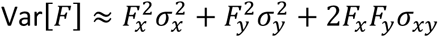

where *F_x_* and *F_y_* denote the partial derivatives of *F* with respect to the variables in the subscript, and 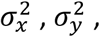, and β_xy_ denote the variances and covariance of *X* and *Y* (Lee et al., 2006). In our case, the random variables are the estimators 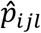 of the connection probabilities which have a variance (squared standard error) of 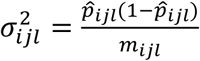 and no covariance because we assume that the two different lineages were measured independently. The denominator*m*_ijl_ denotes the number of tested connections for that particular lineage and combination of layers. This yields the following standard error for *e_ij_*

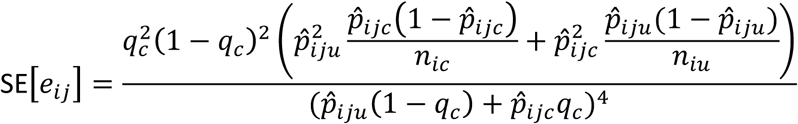

The values reported in Figures 4G and 5E are the estimates and propagated standard errors.

### Log-ratios of connection probabilities between related and unrelated neurons

To visualize the overall pattern of connectivity differences between related and unrelated neurons, we used a heatmap of the log ratio of connection probabilities for related and unrelated neurons (Figure 5F). For each layer-defined connection type, we took the log2 of the ratio of the related pair connection probability and unrelated pair connection probability, with Laplace smoothing (by adding 1 to both the numerator and denominator) applied to both probabilities. Specifically, if *A* out of *B* related pairs and *C* out of *D* unrelated pairs were connected, we computed the log-ratio as log_2_{[(*A*+1)/(*B*+1)] / [(*C*+1)/(*D*+1)]}. The 95% confidence intervals were computed via bootstrapping. For each bootstrap iteration, we generated *A*_boot_ as a binomial draw with *p*=(*A*+1)/(*B*+1) and *n*=*B*, and *C*_boot_ as a binomial draw with *p*=(*C*+1)/(*D*+1) and *n*=*D*. As the 95% confidence interval, we took 95% coverage interval of the bootstrapped log-ratios log_2_{[(*A*_boot_+1)/(*B*+1)] / [(*C*_boot_+1)/(*D*+1)]}.

---

**Figure S1.**
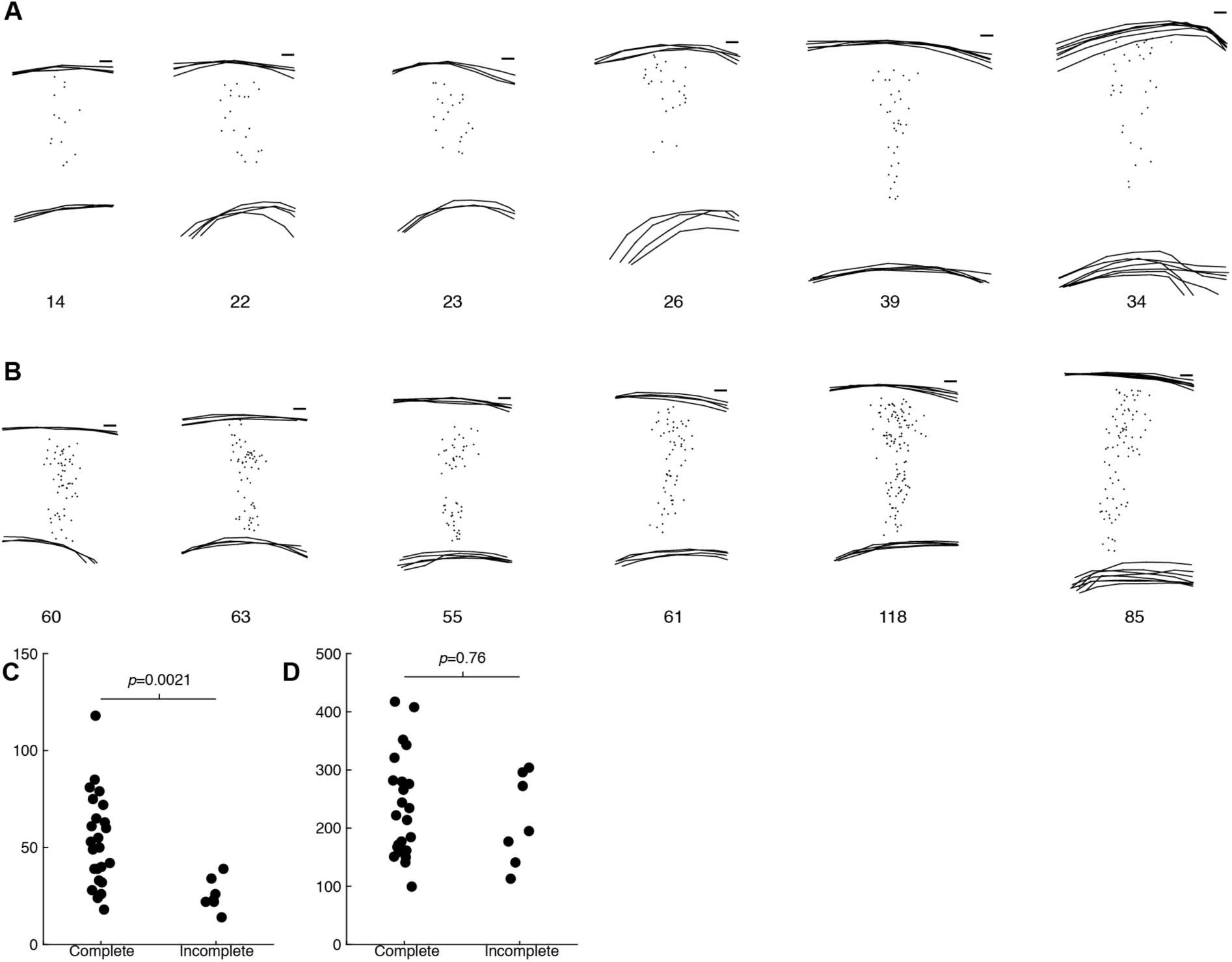
Clones induced at E11.5 are often restricted to superficial layers, related to Figure 1. **(A)** Clones induced at E11.5 that were considered “incomplete” due to lack of neurons in deep cortical layers. **(B)** Examples of clones induced at E11.5 that were considered “complete” due to inclusion of neurons in deep cortical layers. **(C and D)** Incomplete clones contain fewer neurons **(C)** but there is no difference in clone width between complete and incomplete clones (**D**; n = 24 and 7 for complete and incomplete clones; p-values computed using Wilcoxon rank sum test. Scale bars: 100 µm **(A and B)**. Related to Figure 1.

**Figure S2.**
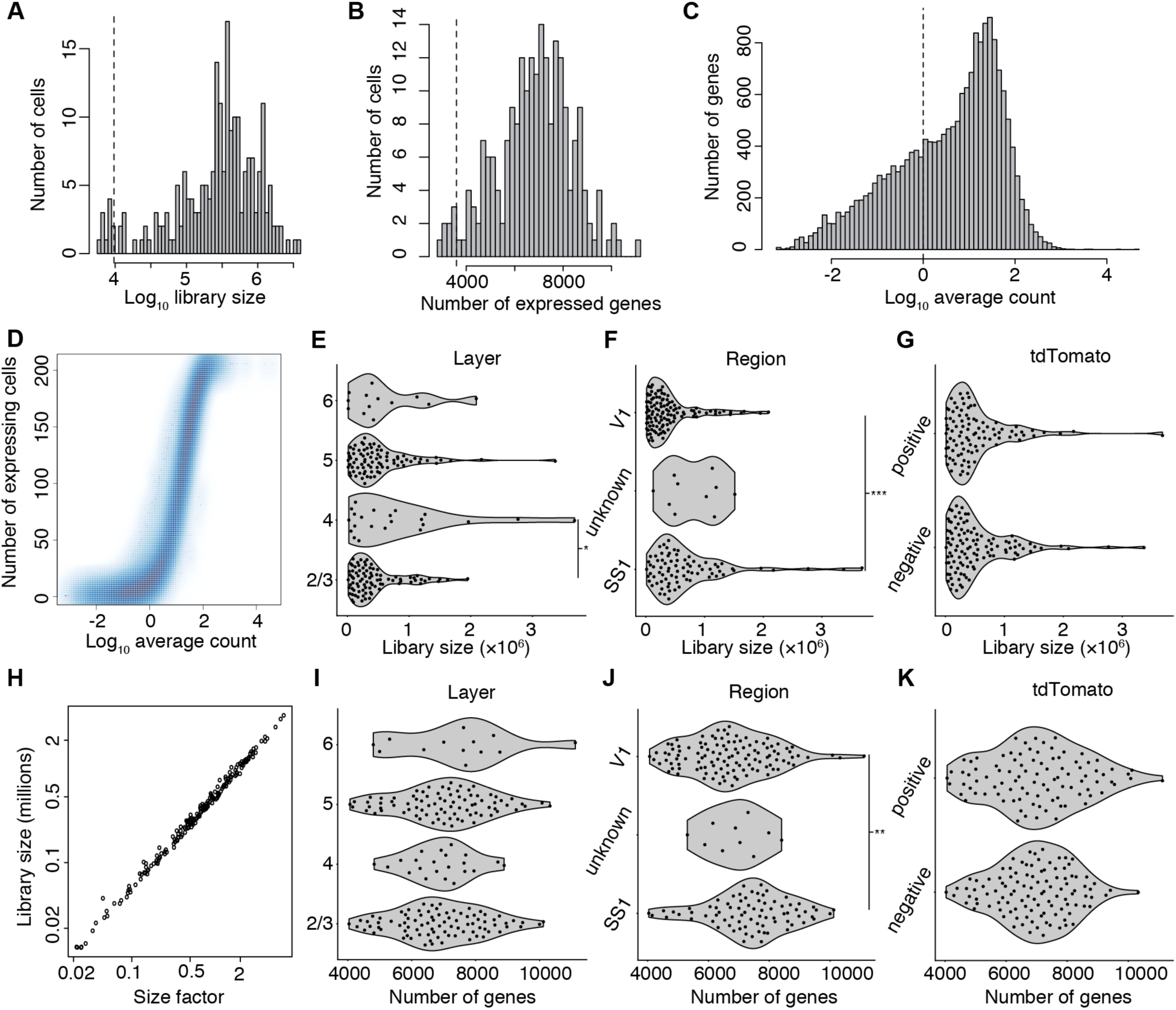
Quality control criteria for single-cell RNA-sequencing data, related to Figure 2. **(A and B)** Histograms of library size **(A)** and number of genes expressed **(B)** for all sequenced cells. Cells falling more than three median absolute differences below the median (dotted lines) were excluded (n=11 based on library size and n=8 based on number of genes expressed; all 8 cells excluded based on number of genes are also excluded based on library size, leaving 206 out of 217 sequenced cells passing these combined criteria). **(C)** Histogram of average number of counts per cell across genes. Genes with less than 1 count per cell on average (dotted line) were excluded from further analyses (n=12,841 genes passing this criteria). **(D)** Correlation between average number of counts per cell and total number of cells expressing each gene, across genes. **(E–G)** Library sizes of cells in different layers (**E**; n=87, 22, 84, and 13 cells in layers 2/3, 4, 5, and 6 respectively; One-way analysis of variance with post-hoc pairwise comparisons using Tukey’s honestly significant difference procedure), cortical regions (**F**; n = 79, 10, and 117 cells in primary somatosensory (SS1), unknown, and primary visual (V1) areas respectively; One-way analysis of variance with post-hoc pairwise comparisons using Tukey’s honestly significant difference procedure), and with (positive) or without (negative) tdTomato-expression (**G**; n = 110 and 96 negative and positive cells respectively; One-way analysis of variance). **(H)** Correlation between size factors used for normalization and library size, across cells. **(I–K)** Number of genes expressed by cells in different layers (**I**; n=87, 22, 84, and 13 cells in layers 2/3, 4, 5, and 6 respectively; One-way analysis of variance), cortical regions (**J;** n = 79, 10, and 117 cells in primary somatosensory (SS1), unknown, and primary visual (V1) areas respectively; One-way analysis of variance with post-hoc pairwise comparisons using Tukey’s honestly significant difference procedure), and with (positive) or without (negative) tdTomato-expression (**K**; n = 110 and 96 negative and positive cells respectively; One-way analysis of variance). Values are raw data points expressed as scatter plots (**D** and **H**, **D** with smoothing), binned **(A–C)**, or with overlay violin plots (**E–G** and **I–K**). Only significant (p<0.05) p-values are shown (*p<0.05; **p<0.01; ***p<0.001). Related to Figure 2.

**Figure S3.**
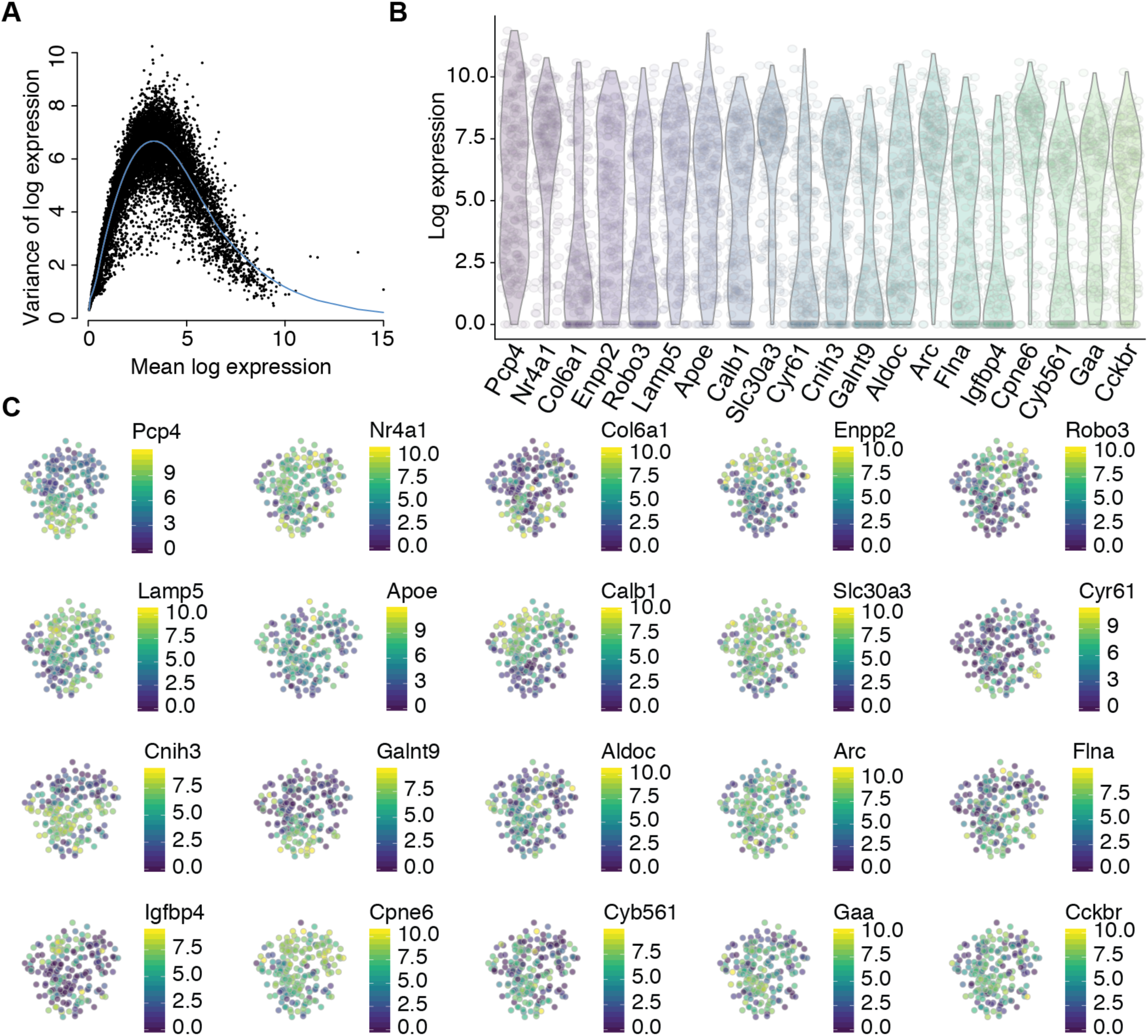
Expression of top highly variable genes, related to Figure 2. **(A)** Variance of normalized log-transformed expression for each gene plotted against the mean log-transformed expression. **(B)** Violin plots of log-transformed expression values for the top twenty highly variable genes across all cells. **(C)** T-distributed stochastic neighbor embedding (t-SNE) was performed using the top highly variable and correlated genes (n=91). Plots are colored by log-transformed expression values for the top twenty highly variable genes. Related to Figure 2.

**Figure S4.**
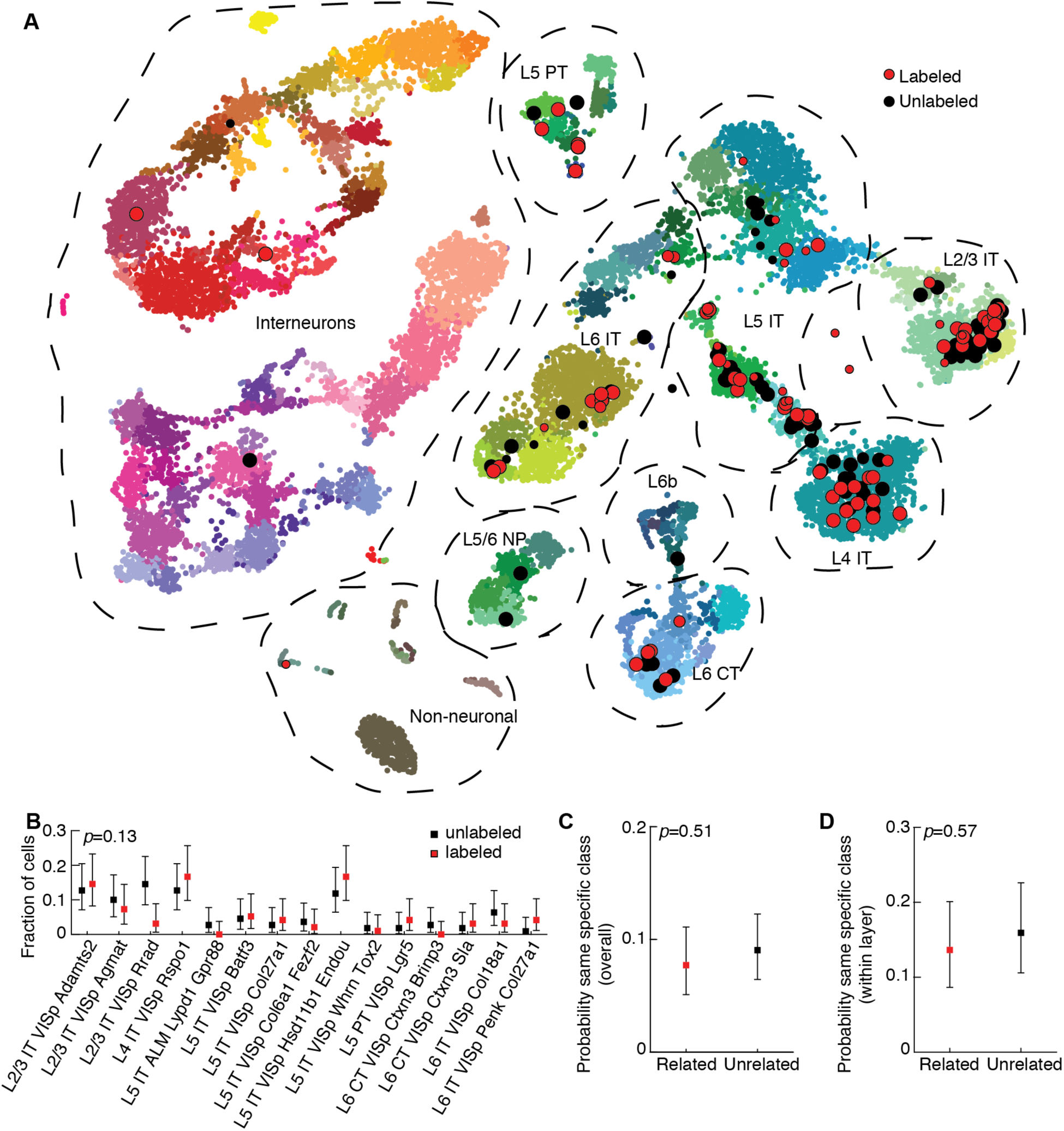
Radial clones are composed of diverse subtypes of excitatory neurons with no evidence of fate restriction, related to Figure 3. **(A)** T-distributed stochastic neighbor embedding (t-SNE) plot similar to **3A**, but with Patch-seq data points (black outline, n=96 and 110 labeled and unlabeled cells respectively) colored according to tdTomato expression status rather than cortical layer, aligned with a recently published mouse cell type atlas (data points with no outline; n=23,822; from (Tasic et al., 2018); colors denote transcriptomic types and are taken from the original publication). The t-SNE of the reference dataset and the positioning of Patch-seq cells were performed as described in (Kobak & Behrens, 2018), see Methods. The size of the Patch-seq data points denotes the precision of the mapping (see Methods): small points indicate high uncertainty. **(B–D)** A similar analysis to **3B–D**, but using specific cell types (tree leaves of the cell type atlas) rather than broad classes. **(B)** Fraction of labeled (n=96) and unlabeled (n=110) cells that mapped to each of the specific clusters identified in (Tasic et al., 2018) with greater than three Patch-seq cells total (no significant difference, Chi-squared test). **(C and D)** Probability of related and unrelated cell pairs mapping to the same specific cluster either overall (**C**; n=337 related pairs, n=409 unrelated pairs; no significant difference, Chi-squared test) or when conditioned on layer position (**D**; n=154 related pairs, n=157 unrelated pairs; no significant difference, Chi-squared test). Related to Figure 3.

**Figure S5.**
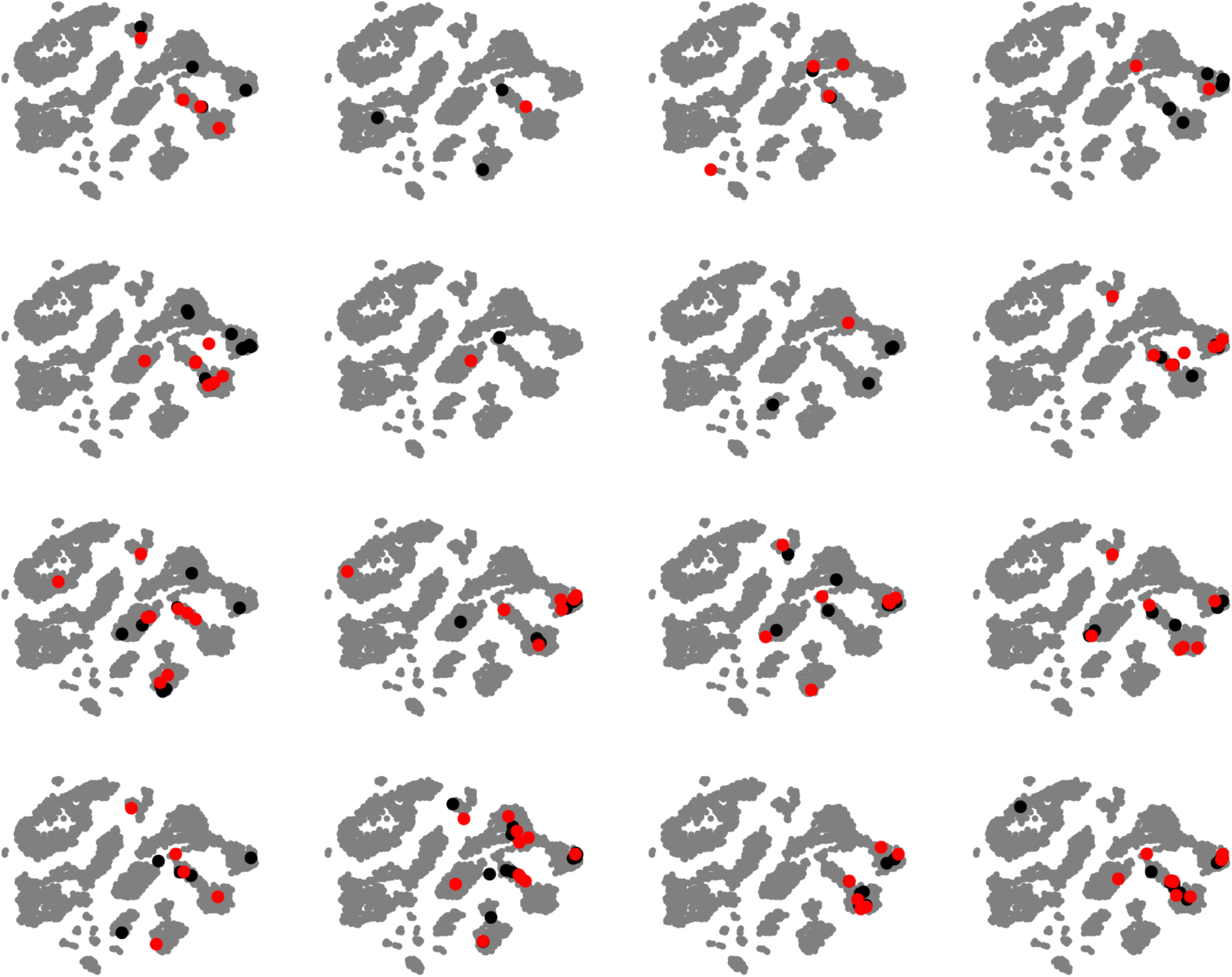
Transcriptomic diversity of individual radial clones, related to Figure 3. t-SNE plots for each Patch-seq experiment (n=16) showing clonally related cells in red and unrelated cells in black, projected onto the reference atlas (grey, from (Tasic et al., 2018)). Related to Figure 3.

**Figure S6.**
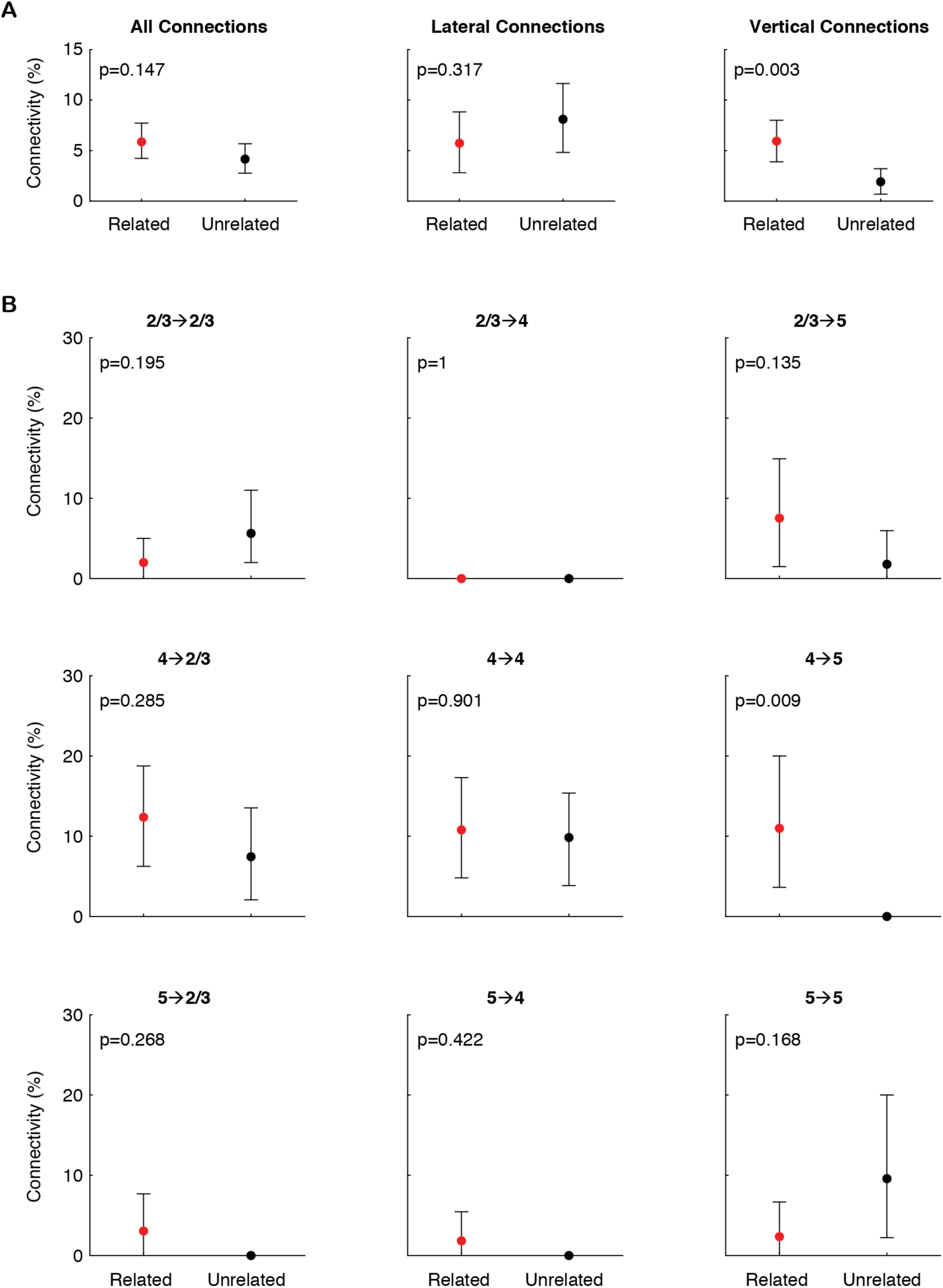
Connectivity differences between clonally related and distance-matched unrelated pairs of neurons, related to Figures 4 and 5. **(A)** Connectivity for all connections (left panel); only lateral, within-layer connections (middle panel); or only vertical, across-layer connections (right panel). **(B)** Connectivity for each layer-defined connection type tested. Pre- and post-synaptic location of the cell bodies is designated above each plot. In **(A and B)**, error bars are 95% coverage intervals computed by resampling (see Methods); *p*-values are two-sided and computed by resampling (see Methods). Related to Figures 4 and 5.

**Figure S7.**
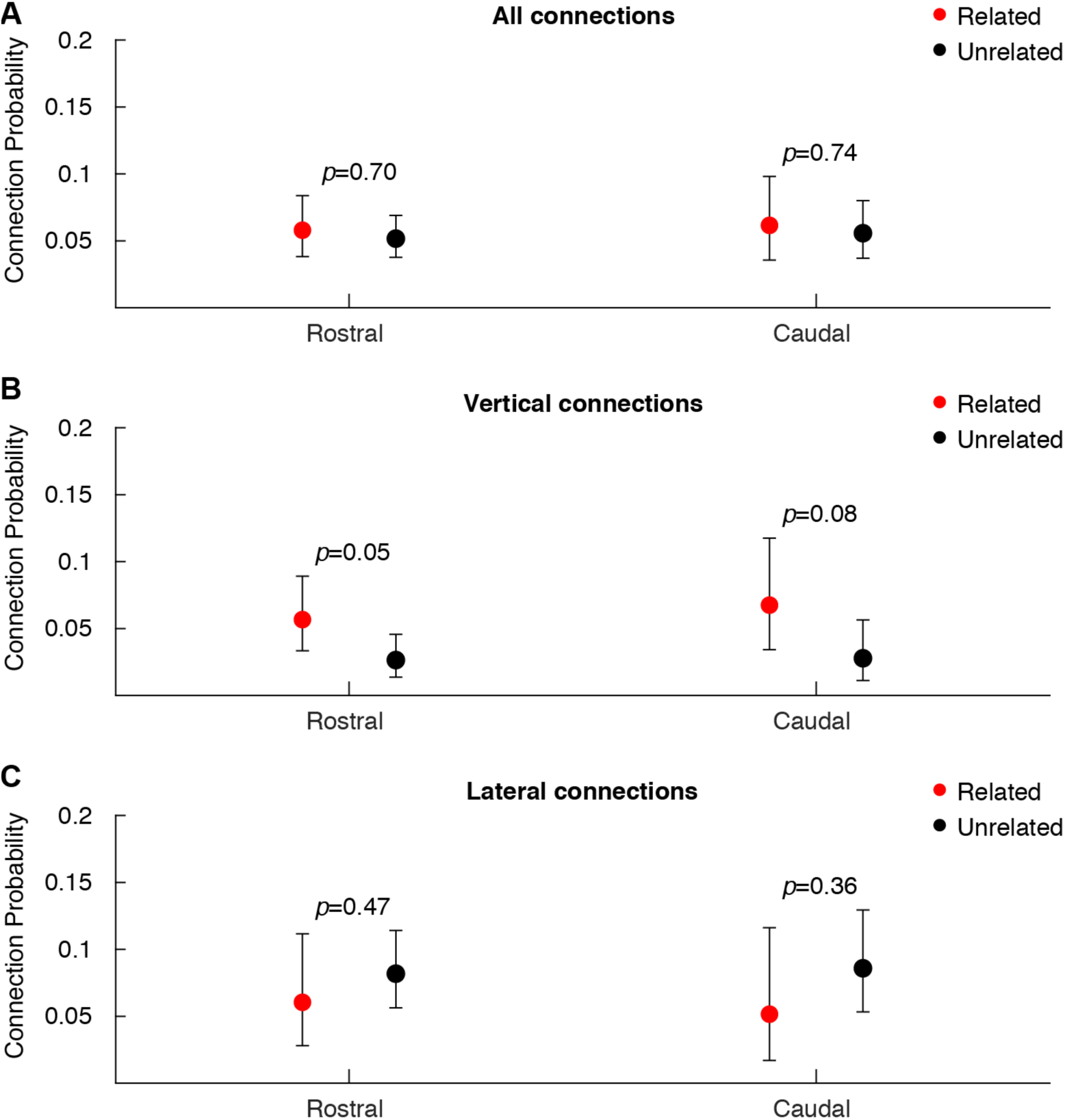
Connectivity differences between clonally related and unrelated neurons across different rostrocaudal location. **(A)** Connection probabilities for all connection types tested, grouped by rostrocaudal position (n=449 and 260 related pairs and n=833 and 485 unrelated pairs in rostral and caudal groups, respectively). **(B)** Connection probabilities for all vertical, across-layer connections tested, grouped by rostrocaudal position (n=300 and 163 related pairs and n=454 and 252 unrelated pairs in rostral and caudal groups, respectively). **(C)** Connection probabilities for all lateral, within-layer connections tested, grouped by rostrocaudal position (n=149 and 97 related pairs and n=379 and 233 unrelated pairs in rostral and caudal groups, respectively). Error bars are 95% Clopper-Pearson confidence intervals and *p*-values are computed using Fisher’s exact test. Related to Figures 4 and 5.

**Table S1 (.xls file). Gene expression data, related to Figure 2.** Normalized counts, normalized log counts, and metadata for all Patch-seq neurons included in our analysis.

**Table S2 (.xls file). Mapping to transcriptomic cell types, related to Figure 3.** Best match for each of our cells onto reference transcriptomic cell types, t-SNE coordinates for the reference dataset, and t-SNE coordinates for projection of our data onto the reference with measure of uncertainty.

**Table S3.** Generalized linear model of connectivity. Connectivity was modeled as a binomial response variable with the following predictors: lineage relationship (1 for related, 0 for unrelated), connection type (1 for vertical, 0 for lateral), Euclidean distance between the cells in microns, and rostrocaudal position (a numeric factor from 1 to 5) (see Methods). ‘×’ denotes an interaction between two linear terms. Overall χ^2^=33.5 compared to constant model, *p*=2.26×10^−4^, 1988 error degrees of freedom. The four terms with small *p*-values are: connection class (connection probability *P* is lower for control vertical connections, compared to control lateral), Euclidean distance (P decreases with increasing distance for unrelated lateral connections), lineage × connection type (*P* is higher for related vertical pairs), and connection type × Euclidean distance (the effect of Euclidean distance on *P* depends on the type of connection tested).

| Term | Estimated<br>Coefficient | SE | t-statistic | p-value |
| --- | --- | --- | --- | --- |
| Constant | -1.82 | 0.43 | -4.22 | $2.37 \cdot 10^{-5}$ |
| Lineage | -0.47 | 0.58 | -0.81 | 0.42 |
| Connection type | -1.55 | 0.75 | -2.06 | 0.039 |
| Euclidean distance | $-8.42 \cdot 10^{-3}$ | $4.40 \cdot 10^{-3}$ | -1.91 | 0.056 |
| Rostrocaudal position | 0.074 | 0.11 | 0.65 | 0.51 |
| Lineage × Connection type | 1.25 | 0.59 | 2.11 | 0.035 |
| Lineage × Euclidean distance | $2.05 \cdot 10^{-4}$ | $2.46 \cdot 10^{-3}$ | 0.083 | 0.93 |
| Lineage × Rostrocaudal position | 0.028 | 0.15 | 0.18 | 0.86 |
| Connection type × Euclidean distance | $7.97 \cdot 10^{-3}$ | $3.99 \cdot 10^{-3}$ | 2.00 | 0.046 |
| Connection type × Rostrocaudal position | $5.86 \cdot 10^{-3}$ | 0.20 | 0.030 | 0.98 |
| Euclidean distance × Rostrocaudal position | $-5.97 \cdot 10^{-4}$ | $9.33 \cdot 10^{-4}$ | -0.64 | 0.52 |

